# Stereotypic Neutralizing V_H_ Clonotypes Against SARS-CoV-2 RBD in COVID-19 Patients and the Healthy Population

**DOI:** 10.1101/2020.06.26.174557

**Authors:** Sang Il Kim, Jinsung Noh, Sujeong Kim, Younggeun Choi, Duck Kyun Yoo, Yonghee Lee, Hyunho Lee, Jongtak Jung, Chang Kyung Kang, Kyoung-Ho Song, Pyoeng Gyun Choe, Hong Bin Kim, Eu Suk Kim, Nam-Joong Kim, Moon-Woo Seong, Wan Beom Park, Myoung-don Oh, Sunghoon Kwon, Junho Chung

**Author notes:** Correspondence to (S.K.); (J.C.). These authors contributed equally to this work (Sang Il Kim, Jinsung Noh).

## Abstract

In six of seven severe acute respiratory syndrome coronavirus 2 (SARS-CoV-2) patients, V_H_ clonotypes, encoded by either immunoglobin heavy variable (IGHV)3-53 or IGHV3-66 and immunoglobin heavy joining (IGHJ)6, were identified in IgG_1_, IgA_1_, and IgA_2_ subtypes, with minimal mutations, and could be paired with diverse light chains, resulting in binding to the SARS-CoV-2 receptor-binding domain (RBD). Because most human antibodies against the RBD neutralized the virus by inhibiting host cell entry, we selected one of these clonotypes and demonstrated that it could potently inhibit viral replication. Interestingly, these V_H_ clonotypes pre-existed in six of 10 healthy individuals, predominantly as IgM isotypes, which could explain the expeditious and stereotypic development of these clonotypes among SARS-CoV-2 patients.

## Main Text

Stereotypic neutralizing antibodies (nAbs) that are identified in convalescent patients can be valuable, providing critical information regarding the epitopes that should be targeted during the development of a vaccine ^1,2^. Those antibodies with naïve sequences, little to no somatic mutations, and IgM or IgD isotypes are precious ^3,4^ because these characteristics effectively exclude the possibility that these nAbs evolved from pre-existing clonotypes that are reactive to similar viruses. This critical phenomenon is referred to as original antigenic sin (OAS), and predisposed antibody-dependent enhancement (ADE) enhancing the severity of viral infections, which can sometimes be fatal, as in the case of the dengue virus vaccine ^5–8^. Several groups have identified nAbs for severe acute respiratory syndrome coronavirus 2 (SARS-CoV-2) ^9–13^, and one report has suggested the possibility that stereotypic nAbs may exist among convalescent patients ^9^. However, stereotypic nAbs for SARS-CoV-2 have not yet been identified. Here, we report stereotypic nAbs for SARS-CoV-2, which were identified by mapping nAbs onto deep immunoglobulin repertoires that were profiled from infected patients. One of these stereotypic nAbs was perfectly naïve and was encoded by immunoglobin heavy variable (IGHV)3-53/IGHV3-66 and immunoglobin heavy joining (IGHJ)6. Furthermore, we also found that these exact V_H_ clonotypes pre-exist in the majority of the healthy population, predominantly as an IgM isotype, which immediately provoked the hypothesis that individuals with this V_H_ clonotype may be able to rapidly evolve potent nAbs and experience favorable clinical features, similar to the human immunodeficiency virus (HIV)-1 response observed among individuals who acquire a unique V_H_ clonotype, featuring a very long heavy chain complementarity determining region (HCDR)3, following exposure to syphilis infection ^14^.

To obtain monoclonal nAbs against SARS-CoV-2, we collected blood samples from seven SARS-CoV-2-infected patients (patients A–G) and used them to generate human antibody libraries. Similar to SARS-CoV, SARS-CoV-2 also uses a spike (S) protein for receptor binding and membrane fusion ^15^. This protein interacts with the cellular receptor angiotensin-converting enzyme II (ACE2) to gain entry into the host cell ^16,17^. A previous report suggested that a human monoclonal antibody, which reacted with the receptor-binding domain (RBD), within the S1 region of the S protein, could hinder the initial interaction between the virus and the cell, effectively neutralizing SARS-CoV-2 ^13^. We confirmed the reactivity of the sera derived from patients against recombinant SARS-CoV-2 S and RBD proteins. Patients A and E, who presented with extensive pneumonic infiltrates, also showed high plasma IgG levels against all recombinant SARS-CoV-2 nucleocapsid (NP), S, S1, S2, and RBD proteins, which could be detected 11, 17, and 45 days after symptom onset in Patient A and 23, 44, and 99 days after symptom onset in Patient E (Supplementary Table 1 and Supplementary Fig. 1). Notably, the sera samples from Middle East respiratory syndrome coronavirus (MERS-CoV) patients cross-reacted with the SARS-CoV-2 S protein, showing a higher titer against the S2 domain, and vice versa (Supplementary Fig. 1 and 2), suggesting the potential risk for ADE. We generated four human antibody libraries, utilizing a phage display system, based on the blood samples from Patient A, which were collected on days 17 and 45 (A_d17 and A_d45), and Patient E, which were collected on days 23 and 44 (E_d23 and E_d44). After biopanning, we successfully isolated 38 single-chain variable fragment (scFv) clones that were reactive against recombinant SARS-CoV-2 RBD in an enzyme immunoassay (Supplementary Fig. 3 and Supplementary Table 2). The half-maximal binding of these scFv-human kappa light chain fragment (hCκ) fusion proteins with the coated antigens occurred at concentrations ranging from 0.32 to 364 nM, which was compatible with the findings of previous reports that have described human monoclonal antibodies against SARS-CoV-2 RBD ^10,13^. Then, we tested whether these antibody clones could inhibit the binding between recombinant SARS-CoV-2 S protein and Vero E6 cells expressing the ACE2 receptor. When incubated with 1.5 × 10^5^ Vero E6 cells, the recombinant HIS-tagged SARS-CoV-2 S protein showed saturated binding at 200 nM, according to flow cytometry analysis, using a fluorescein isothiocyanate (FITC)-labeled anti-HIS antibody. For the analysis, recombinant S protein (200 nM) was mixed with scFv-hFc fusion proteins, at a final concentration of either 200 nM (equimolar) or 600 nM (molar ratio of 1:3). Eleven clones (A-1A1, A-1H4, A-1H12, A-2F1, A-2H4, E-2G3, E-3A12, E-3B1, E-3G9, E-3H31, and E-4D12) almost completely inhibited the binding between recombinant S protein and Vero E6 cells at 600 nM, and some showed potent inhibition activity, even at 200 nM (Supplementary Fig. 4). The neutralizing potency of these 11 clones for the inhibition of viral replication was tested using an *in vitro* assay. Vero cells, in a T-25 flask, were infected with SARS-CoV-2, at a medium tissue culture infectious dose (TCID_50_) of 2,500 and in the presence of scFv-hCκ fusion proteins, at concentrations of 0.5, 5, or 50 μg/mL. Viral RNA concentrations in the culture supernatant were determined 0, 24, 48, and 72 hours after infection. Nine antibodies exhibited complete neutralizing activity, at 50 μg/mL (Supplementary Fig. 5), and two antibodies (A-1H4 and E-3G9) showed potent neutralization, even at 5 μg/mL (Supplementary Fig. 5).

We also performed deep profiling of the immunoglobulin (IG) repertoire in three chronological blood samples each from patients A and E and two chronological samples from each of the other five patients. Then, we searched for nAb clonotypes that possessed identical VJ combinations and perfectly matched HCDR3 sequences, at the amino acid level among the immunoglobulin heavy chain (IGH) repertoires of Patients A and E. One and five nAb clonotypes were successfully identified in Patients A and E, respectively (Fig. 1a). Notably, three nAbs (A-2F1, E-3A12, and E-3B1) were encoded by IGHV3-53/IGHV3-66 and IGHJ6 (Fig. 1a). These two V_H_ genes, IGHV3-53*01 and IGHV3-66*01, are identical at the amino acid level, except for the H12 residue (isoleucine in IGHV3-53 and valine in IGHV3-66), and only five nucleotide differences exist between their sequences. Furthermore, four clonotypes were IgG_1_, and two clonotypes class-switched to IgA_1_ and IgA_2_ when examined 44 days after symptom onset (Fig. 1a). These clonotypes had a very low frequency of somatic mutations (1.03% ± 0.51%), which was compatible with findings regarding other nAbs in previous reports ^9,10^. Then, we collected all V_H_ sequences from the seven patients and searched the clonotypes of 11 nAbs that were encoded by the same V_H_ and J_H_ genes and showed 66.6% or higher identity in the HCDR3 sequence, at the amino acid level (Supplementary Fig. 6). Interestingly, clonotypes that were highly homologous to the E-3B1 nAb were found among six of seven patients, with a total of 55 sequences among the isotypes IgG_1_ (Patient A, B, D, E, F, and G), IgA_1_ (Patient E and G), and IgA_2_ (Patient E) (Supplementary Table 3). These clonotypes shared nearly identical V_H_ sequences (92.78% ±1.40% identity at the amino acid level), with E-3B1 displaying an extremely low frequency of somatic mutations (0.77% ± 0.93%). Among these 55 clonotypes, 22 unique HCDR3s were identified, at the amino acid level, and eight unique HCDR3s existed in more than one patient. To test the reactivity of clonotypes homologous to E-3B1 against the SARS-CoV-2 S protein, we arbitrarily sampled 12 IGH clonotypes (Fig. 1b), containing five different HCDR3s, from the IGH repertoires of six patients. The genes encoding these IGH clonotypes were chemically synthesized and used to construct scFv genes, using the V_λ_ gene from the E-3B1 clone. Then, the reactivities of these scFv clones were tested in an enzyme immunoassay. Three clones (E-12, A-32, and B-33) reacted against the recombinant S and RBD proteins (Fig. 1b). Then, scFv libraries were constructed, using the A-11, A-31, E-34, A,B,G-42, G-44, D-51, F-53, E-52, and A-54 genes, and the V_κ_/V_λ_ genes were amplified from Patients A, E, and G. Consequently, we confirmed that all 12 IGH clonotypes were reactive against both recombinant S and RBD proteins when paired with eight different V_κ_ and V_λ_ genes (Fig. 1b,c). Moreover, all seven patients possessed these V_κ_/V_λ_ clonotypes with identical VJ gene usage and perfectly matched LCDR3 amino acid sequences (Supplementary Fig. 7). In particular, IGLV2-14/IGLJ3, IGLV3-19/IGLJ2, and IGLV3-21/IGLJ2 were frequently used across all seven patients (Supplementary Fig. 8 and 9). Because E-3B1 effectively inhibited the replication of SARS-CoV-2 (Fig. 1d), these 55 clonotypes are likely to neutralize the virus when paired with an optimal light chain.

**Fig. 1.**
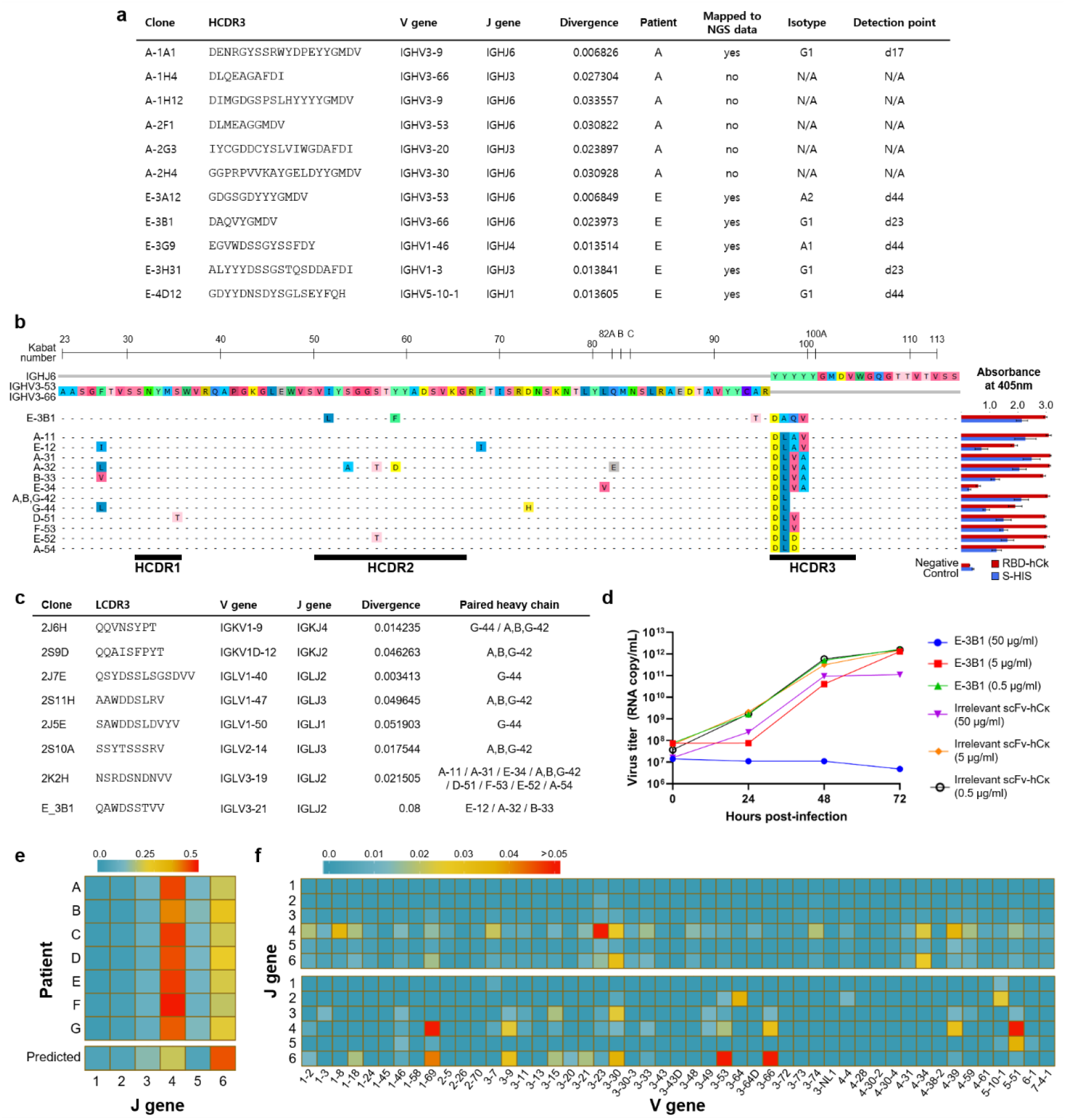
Characteristics of nAbs, derived from patients A and E, stereotypic IGH clonotypes that are highly homologous to E-3B1, and the predicted RBD-binding clones that were enriched through biopanning. Stereotypic nAb V_H_ clonotypes against the SARS-CoV-2 RBD, encoded by IGHV3-53/3-66 and IGHJ6, were found in six of seven patients. **a,** Characteristics of nAbs discovered in patients A and E. **b,** IGH clonotypes that are highly homologous to E-3B1 and reactive against recombinant SARS-CoV-2 S and RBD proteins. The right column shows the results of the phage ELISA. All experiments were performed in quadruplicate, and the data are presented as the mean ± SD. **c,** List of diverse IGL clonotypes that can be paired with the IGH clonotypes from **b** to achieve reactivity. **d,** Measurement of viral RNA in the culture supernatant of Vero cells after SARS-CoV-2 infection **e,** J and **f,** VJ gene usage in the IGH repertoire of patients (upper) and the binding-predicted IGH clones (bottom). For the VJ gene usage heatmap, the frequency values for the IGH repertoire of all seven patients were averaged and are displayed (upper) along with those of the predicted RBD-binding IGH clones (bottom). N/A: not applicable

Among these IGH clonotypes, A,B,G-42 was quite unique, presenting no somatic mutations and containing an HCDR3 (DLYYYGMDV) formed by the simple joining of IGHV3-53 and IGHJ6. This naïve V_H_ sequence existed in the IGH repertoire of three patients (A, B, and G), as IgG_1_, IgG_1_, or IgG_1_ and IgA_1_ subtypes, respectively (Table 1). More interestingly, the IGH clonotypes encoded by IGHV3-53/IGHV3-66 and IGHJ6 that possessed an HCDR3 (DLYYYGMDV) with zero to one somatic mutation residues could be identified within the IGH repertoire of six of 10 healthy individuals, predominantly as an IgM isotype (Table 1), based on publicly available IGH repertoires ^18^. The A,B,G-42 clonotype showed light chain plasticity and paired with five V_κ_ /V_λ_ genes to achieve RBD binding. In particular, the V_κ_ gene (2J6H) accumulated only five somatic mutations (1.4% divergence). None of the 12 clones, including A,B,G-42, reacted against the recombinant RBD proteins from either SARS-CoV or MERS-CoV (Supplementary Fig. 10). In our prior experiment, none of the 37 identified MERS-RBD-binding human monoclonal antibodies, from two patients, were encoded by IGHV3-53/IGHV3-66 and IGHJ6 (Supplementary Table 4) ^19^. Therefore, the presence of these stereotypic-naïve IGH clonotypes in the healthy population, and their light chain plasticity to achieve SARS-CoV-2 RBD binding, may be unique to SARS-CoV-2, which might provide a rapid and effective humoral response to the virus among patients who express these clonotypes. These findings provide the majority of the population possess germline-precursor B cells, encoded by IGHV3-53/IGHV3-66 and IGHJ6, which can actively initiate virus neutralization upon SARS-CoV-2 infection.

**Table 1.**
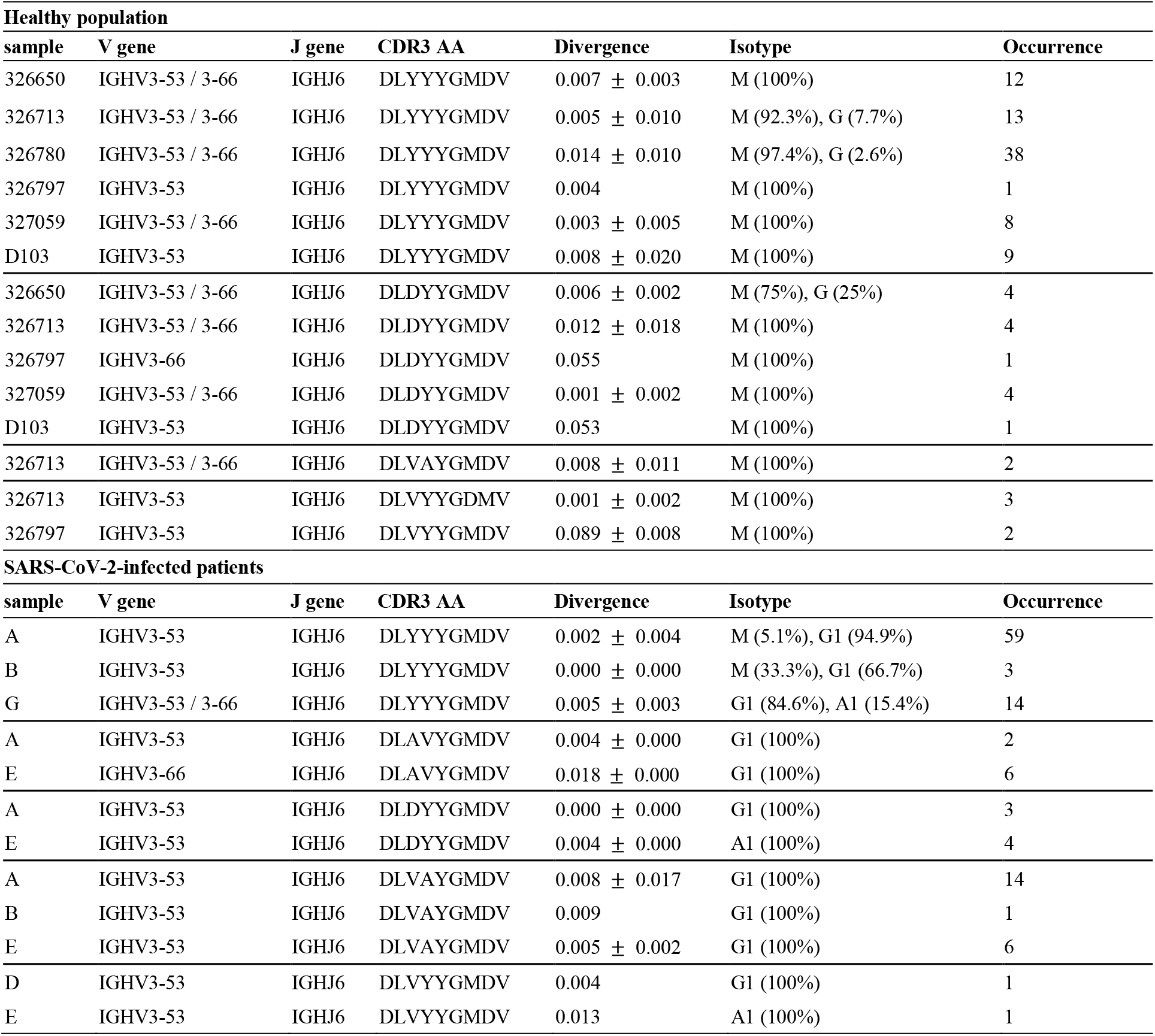
The stereotypic V_H_ clonotypes against SARS-CoV-2 RBD in the healthy population and patients. The healthy samples based on publicly available IGH repertoires or patient identification can be found in the sample column. Clonotypes were mapped according to identical VJ gene usage of IGHV3-53/IGHV3-66 and IGHJ6 and perfectly matched HCDR3 at the amino acid level. The read counts of the mapped sequences in the repertories of each samples were annotated in the occurrence column. For the clonotypes with multiple occurrences, mean and standard deviation of divergence were represented. The proportion of each isotypes were indicated for the all samples.

To further elucidate the preferential use of IGHV3-53/IGHV3-66 and IGHJ6 genes during the generation of SARS-CoV-2 RBD-binding antibodies, we extracted 252 predicted RBD-binding clones from our biopanning data (See Methods). We previously showed that antibody clones with binding properties can be predicted by employing next-generation sequencing (NGS) technology and analyzing the enrichment patterns of biopanned clones ^20,21^. Although the IGHJ4 gene was more prominent in the IGH repertoires of the seven patients, similar to healthy human samples ^18,22^, the predicted RBD-binding clones primarily used the IGHJ6 gene (Fig. 1e). Furthermore, the predicted RBD-binding clones showed the dominant usage of IGHV3-53/IGHJ6 and IGHV3-66/IGHJ6 pairs, which was not observed in the whole IGH repertoires of patients (Fig. 1f).

Naïve B cells typically undergo somatic hypermutations, clonal selection, and class-switching following antigen exposure. We examined the chronological events that occurred in all IGH clonotypes identified in patients and those that were reactive against the SARS-CoV-2 RBD. We categorized RBD-reactive clones into three groups: neutralizing antibodies (neutralize), binding-confirmed antibodies (bind), and binding-predicted antibodies (predicted). In all three groups, these IGH clonotypes appeared and disappeared along the disease course and showed a low frequency of somatic mutations (Fig. 2c,d) and rapid class-switching, especially to IgG_1_, IgA_1_, and IgA_2_. In the entire IGH repertoire of the patients, naïve-derived IGH clonotypes with minimal somatic mutations (< 2.695% ± 0.700%) showed increased IgG_3_ and IgG_1_ subtypes, and the proportion of IgG_1_ subtype was dramatically increased for a period (Fig. 2a,b and Supplementary Fig. 11). Furthermore, these naïve-derived IGH clonotypes were detected as IgA_1_ and IgG_2_ subtypes in patients A and E, as minor populations (Fig. 2a,b), and as the IgA_2_ subtype in Patient E (Fig. 2b). To summarize, RBD-reactive IGH clonotypes rapidly emerged and underwent class-switching, to IgG_1_, IgA_1_, and IgA_2_, without experiencing many somatic mutations. However, this dramatic temporal surge of naïve IGH clonotypes, with rapid class-switching, occurred across the entire IGH repertoire of patients and was not confined to those reactive to the SARS-CoV-2 RBD. Because several mutations within the RBD have been identified along the course of the SARS-CoV-2 pandemic, worldwide ^23^, we examined the probability of emerging escape mutants from the IGH repertoire induced by the wild-type virus infection. Our E-3B1, A-1H4, A-2F1, A-2H4, and E-3G9 nAbs successfully bound to recombinant mutant RBD proteins (V341I, F342L, N354D/D364Y, V367F, A435S, W436R, G476S, and V483A) in a dose-dependent manner, with compatible reactivity against recombinant wild-type RBD protein (Supplementary Fig. 12). Therefore, the human IGH immune repertoire may provide effective protection against most current SARS-CoV-2 mutants.

**Fig. 2.**
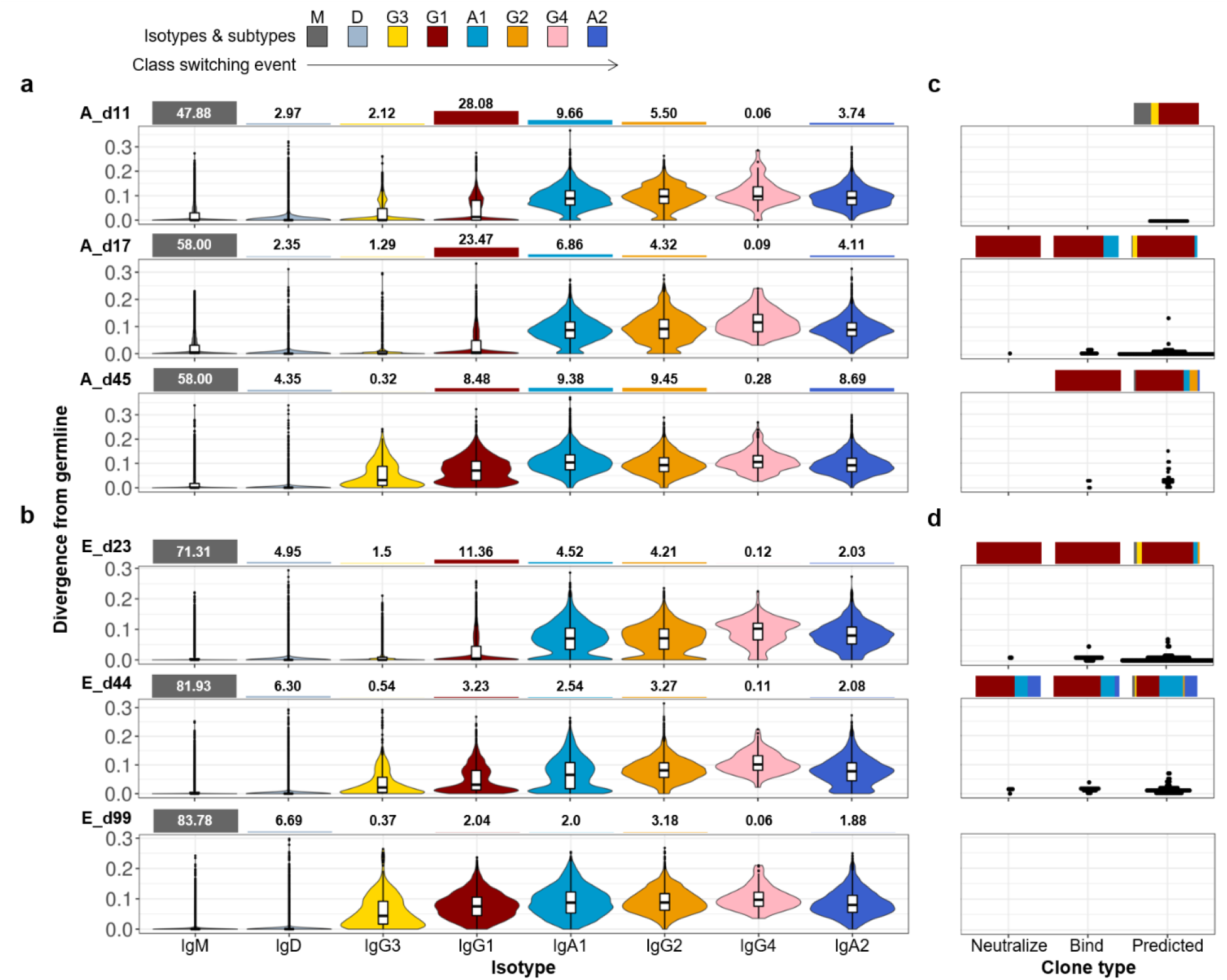
Deep profiling of the IGH repertoires of patients A and E. RBD-reactive IGH clonotypes rapidly undergo class-switching events to IgG_1_, IgA_1_, and IgA_2_, with few somatic mutations. **(a,b)** IGH repertoires of **a,** Patient A and **b,** Patient E were analyzed 11, 17, and 45 (A_d11, A_d17, A_d45) days and 23, 44, and 99 (E_d23, E_d44, E_d99) days after symptom onset, respectively. IGH repertoires were examined according to divergence from the germline and the isotype composition of the sequences. Values for divergence from the germline were calculated separately for each isotype and are presented as violin plots, ordered by the class-switch event. The bar graphs on the top of the violin plots represent the proportion of each isotype in the repertoire. (**c,d)** Mapping of three types of RBD-binding IGH sequences (neutralize, bind, and predicted), derived from either **c,** Patient A or **d,** Patient E, against the corresponding IGH repertoire. The positions of the RBD-binding IGH sequences in the divergence value were annotated as dot plots, on the same scale used for **a** and **b**. Bar graphs on the top of the dot plots indicate the isotype compositions of the sequences in the repertoire.

In response to SARS-CoV-2 infection, most human IGH repertoires efficiently generated clonotypes encoded by IGHV3-53/IGHV3-66 and IGHJ6, which could be paired with diverse light chains, for both RBD binding and virus neutralization, with few to no somatic mutations. These clonotypes underwent swift class-switching to IgG_1_, IgA_1_, and even IgA_2_ subtypes. The expeditious development of these IGH clonotypes would be possible because the naïve-stereotypic IGHV3-53/IGHV3-66 and IGHJ6 clonotypes pre-exist in the majority of the healthy population, predominantly as an IgM isotype. In line with our findings, several groups have previously reported potent human nAbs, composed of either IGHV3-53 or IGHV3-66 and IGHJ6 genes, using single B cell sequencing technology ^9–13^. Furthermore, the crystal structures of two IHGV3-53 neutralizing antibodies were determined which showed that two key motifs within HCDR1 and HCDR2 encoded in the IGHV3-53 germline are making contact with RBD ^24^. Therefore, the preferential use of IGHV3-53/IGHV3-66 and IGHJ6 in the development of nAbs to SARS-CoV-2 appeared prominent. From these observations, we hypothesize that the existence of this unique, naïve IGH clonotype would provide near-immediate protection to some people exposed to SARS-CoV-2, and a very favorable clinical course, unlike SARS-CoV or MERS-CoV. In addition, the chronological follow-up of IGH clonotypes, encoded by the IGHV3-53/IGHV3-66 and IGHJ6 genes, along with their class-switching events, would be valuable for the development of a safe and effective vaccine.

## Methods

### Human samples

Three chronological blood samples were drawn from Patients A and E. From Patients B, C, D, F, and G, two chronological samples were obtained. All patients were confirmed to be infected by SARS-CoV-2 by a positive reverse transcriptase-quantitative polymerase chain reaction (RT-qPCR) result, and sample collection was performed at Seoul National University Hospital. Peripheral blood mononuclear cells (PBMCs) and plasma were isolated using Lymphoprep (Stemcell Technologies, Vancouver, BC, Canada), according to the manufacturer’s protocol. The PBMCs were subjected to total RNA isolation, using the TRI Reagent (Invitrogen, Carlsbad, CA, USA), according to the manufacturer’s protocol. The study involving human sample collection was approved by the Institutional Ethics Review Board of Seoul National University Hospital (IRB approval number: 2004-230-1119).

### Next-generation sequencing (NGS)

Genes encoding V_H_ and part of the CH1 domain were amplified, using specific primers, as described previously ^18,25^. All primers used are listed in Supplementary Table 8. Briefly, total RNA was used as a template to synthesize cDNA, using the Superscript IV First-Strand Synthesis System (Invitrogen), with specific primers targeting the constant region (CH1 domain) of each isotype (IgM, IgD, IgG, IgA, and IgE) ^25^, according to the manufacturer’s protocol. Following cDNA synthesis, 1.8 volumes of SPRI beads (AmpureXP, Beckman Coulter, Brea, CA, USA) were used to purify cDNA, which was eluted in 40 μ L water. The purified cDNA (18 μ L) was subjected to second-strand synthesis in a 25-μ L reaction volume, using V gene-specific primers ^18^ and KAPA Biosystems (KAPA HiFi HotStart, Roche, Basel, Switzerland). The PCR conditions were as follows: 95°C for 3 min, 98°C for 1 min, 55°C for 1 min, and 72°C for 5 min. Following the second-strand synthesis, double-strand DNA (dsDNA) was purified, using SPRI beads, as described above. V_H_ genes were amplified using 15 μ L eluted dsDNA and 2.5 pmol of the primers listed in Supplementary Supplementary Table 8, in a 50-μL total reaction volume (KAPA Biosystems), using the following thermal cycling program: 95°C for 3 min; 17 cycles of 98°C for 30 sec, 65°C for 30 sec, and 72°C for 1 min 10 sec; and 72°C for 5 min. The number of PCR cycles was increased, from 17 to 19, for samples from Patients B (d10 and 19), C (d6), E (d23), and G (d9 and 22). PCR products were purified using SPRI beads and eluted in 30 μL water. Genes encoding V_κ_ and V_λ_ were amplified using specific primers, as described previously ^22,26^. Briefly, total RNA was used as a template to synthesize cDNA, using the Superscript IV First-Strand Synthesis System (Invitrogen), with specific primers targeting the constant region, which are listed in Supplementary Table 8, according to the manufacturer’s protocol. Following cDNA synthesis, SPRI beads were used to purify cDNA, which was eluted in 40 μL water. Purified cDNA (18 μ L) was used for the first amplification, in a 25-μ L reaction volume, using VJ gene-specific primers, which are listed in Supplementary Table 8, and KAPA Biosystems. The PCR conditions were as follows: 95°C for 3 min, 4 cycles of 98°C for 1 min, 55°C for 1 min, and 72°C for 1 min; and 72°C for 10 min. Subsequently, DNA was purified using SPRI beads, and the V_κ_ and V_λ_ genes were amplified using 15 μL eluted dsDNA and 2.5 pmol of the primers listed in Supplementary Table 8, in a 50-μL total reaction volume (KAPA Biosystems). The PCR conditions were as follows: 95°C for 3 min; 17 cycles of 98°C for 30 s ec, 65°C for 30 sec, and 72°C for 1 min 10 sec; and 72°C for 5 min. PCR products were purified using SPRI beads, as described above. For the amplification of V_H_ from each round of biopanning (rounds 0–4), gene fragments were amplified from phagemid DNA, using the primers listed in Supplementary Table 8. SPRI-purified sequencing libraries were quantified with a 4200 TapeStation System (Agilent Technologies), using a D1000 ScreenTape Assay, before performing sequencing on an Illumina MiSeq Platform.

## NGS data processing

### Pre-processing of the NGS data for the IG repertoire

The raw NGS forward (R1) and reverse (R2) reads were merged by PEAR, v0.9.10, in default setting ^27^. The merged reads were q-filtered using the condition q20p95, which results in 95% of the base-pairs in a read having Phread scores higher than 20. The location of the primers was recognized from the q-filtered reads while allowing one substitution or deletion (Supplementary Table 8). Then, primer regions that specifically bind to the molecules were trimmed in the reads, to eliminate the effects of primer synthesis errors. Based on the primer recognition results, unique molecular identifier (UMI) sequences were extracted, and the reads were clustered according to the UMI sequences. To eliminate the possibility that the same UMI sequences might be used for different read amplifications, the clustered reads were sub-clustered, according to the similarity of the reads (Five mismatches were allowed in each sub-cluster). The sub-clustered reads were aligned, using a multiple sequence alignment tool, Clustal Omega, v1.2.4, in default setting ^28,29^. From the aligned reads, the frequency of each nucleotide was calculated, and a consensus sequence of each sub-cluster was defined using the frequency information. Then, the read count of the consensus sequence was re-defined as the number of UMI sub-clusters that belong to the consensus sequences.

### Sequence annotation, functionality filtering, and throughput adjustment

Sequence annotation consisted of two parts, isotype annotation and VDJ annotation. For annotation, the consensus sequence was divided into two sections, a VDJ region and a constant region, in a location-based manner. For isotype annotation, the extracted constant region was aligned with the IMGT (international immunogenetics information system) constant gene database ^30^. Based on the alignment results, the isotypes of the consensus sequences were annotated. Then, the VDJ regions of the consensus sequences were annotated, using IgBLAST, v1.8.0 ^31^. Among the annotation results, V/D/J genes (V/J genes for V_L_), CDR1/2/3 sequences, and the number of mutations from the corresponding V genes were extracted, for further analysis. Divergence values were defined as the number of mutations identified in the aligned V gene, divided by the aligned length. Then, the non-functional consensus reads were defined using the following criteria and filtered-out: 1. sequence length shorter than 250 bp; 2. existence of stop-codon or frame-shift in the full amino acid sequence; 3. annotation failure in one or more of the CDR1/2/3 regions; and 4. isotype annotation failure. Then, the functional consensus reads were random-sampled, to adjust the throughput of the V_H_ data (Supplementary Table 5). Throughput adjustment was not conducted for V_L_ data (Supplementary Table 6).

### Pre-processing of the biopanning NGS data

Pre-processing of the biopanning NGS data was performed as previously reported, except for the application of the q-filtering condition q20p95 instead of q20p100 ^32^.

### Overlapping IGH repertoire construction

To investigate the shared IGH sequences among the patients, we defined the overlapping IGH repertoire of the patients. First, histograms for the nearest-neighbor distances of the HCDR3 amino acid sequences were calculated for the repertoire data. A hierarchical, distance-based analysis, which was reported previously ^33^, was applied to the HCDR3 amino acid sequences, to cluster the IGH sequences at a functional level. The IGH sequences for all repertoire data could be approximated into a bimodal distribution, allowing the functionally similar IGH sequences to be extracted by capturing the first peak of the distribution (Supplementary Fig. 13). Threshold values for each data set were defined as the nearest-neighbor distance value of those points with a minimum frequency between the two peaks of the distribution. Then, the minimum value among all threshold values, 0.113871, was used to construct the overlapping IGH repertoire, which means that 11.3871% of mismatches in the CDR3 amino acid sequence were allowed in the overlapping IGH repertoire construction. To construct the overlapping IGH repertoire, the repertoire data sets of all patients were merged into one data set. The IGH sequences in the merged data set were then clustered, using the following conditions: 1. the same V and J gene usage; and 2. mismatch smaller than 11.3871% among the CDR3 amino acid sequences. Subsequently, clusters containing IGH sequences from more than one patient were included in the overlapping IGH repertoire data set.

### Extraction of binding-predicted clones

From each round of biopanning (rounds 0, 2, 3, and 4), the V_H_ genes were amplified and subjected to NGS analysis, using the MiSeq platform, as described previously ^21^. Binding-predicted clones from biopanning were defined by employing frequency the values of the NGS data from four libraries, A_d17, A_d45, E_d23, and E_d44, at each round of biopanning. The enrichment of clones primarily occurred during the second round of biopanning, based on the input/output virus titer values for each round of biopanning and the frequencies of the clones in the NGS data (Supplementary Fig. 14). Then, the frequency information in the NGS data sets for biopanning rounds 0, 2, 3, and 4 was subject to principal component analysis (PCA), for dimension reduction. Accordingly, principal component (PC)1 and PC2, which represented clone enrichment and clone depletion, respectively, were extracted. In the biopanning data, PC1 was primarily composed of the frequencies in rounds 2, 3, and 4, whereas PC2 was primarily composed of the frequency in round 0 (Supplementary Fig. 15). Thus, we defined PC1-major clones as the predicted clones, by setting constant threshold values on the PC1 value and the ratio between PC1 and PC2 (Supplementary Table 7). Subsequently, 94.74% of the RBD-binding clones were successfully mapped to the predicted clones (Supplementary Fig. 15).

### Construction of a human scFv phage-display library and V_L_ shuffled libraries

For the V_H_ gene, the cDNA prepared for the NGS analysis was used. For the V_Κ_ and V_λ_ genes, total RNA was used to synthesize cDNA, using the Superscript IV First-Strand Synthesis System (Invitrogen), with oligo(dT) primers, according to the manufacturer’s instructions. Then, the genes encoding V_Κ_/V_λ_ and V_H_ were amplified, from the oligo(dT)-synthesized cDNA and the cDNA prepared for NGS analysis, respectively, using the primers listed in Supplementary Table 8 and KAPA Biosystems. The PCR conditions were as follows: preliminary denaturation at 95°C for 3 min; 4 cycles of 98°C for 1 min, 55°C for 1 min, and 72°C for 1 min; and 72°C for 10 min. Subsequently, DNA was purified using SPRI beads, as described above. The purified DNA was amplified using the primers listed in Supplementary Table 8 and KAPA Biosystems. The PCR conditions were as follows: preliminary denaturation, at 95°C for 3 min; 25 cycles of 98°C for 30 sec, 58°C for 30 sec, and 72°C for 90 sec; and 72°C for 10 min. Then, the V_H_ and V_Κ_/V_λ_ fragments were subjected to electrophoresis, on a 1% agarose gel, and purified, using a QIAquick Gel Extraction Kit (Qiagen Inc., Valencia, CA, USA), according to the manufacturer’s instructions. The purified V_H_ and V_Κ_/V_λ_ fragments were mixed, at equal ratios at 50 ng, and subjected to overlap extension, to generate scFv genes, using the primers listed in Supplementary Table 8 and KAPA Biosystems. The PCR conditions were as follows: preliminary denaturation, at 94°C for 5 min; 25 cycles of 98°C for 15 sec, 56°C for 15 sec, and 72°C for 2 min; and 72°C for 10 min. The amplified scFv fragment was purified and cloned into a phagemid vector, as described previously ^34^.

For the construction of V_Κ_/V_λ_ shuffled libraries, gBlocks Gene Fragments (Integrated DNA Technologies, Coralville, IA, USA), encoding A-11, E-12, A-31, A-32, B-33, E-34, A,B,G-42, G-44, D-51, F-53, E-52, and A-54, were synthesized. Synthesized V_H_ and the V_Κ_/V_λ_ genes from Patients A, E, and G were used to synthesize the scFv libraries using PCR, as described previously ^34^. Then, the amplified scFv fragments were purified and cloned into the phagemid vector, as described above.

### Biopanning

A phage display of the human scFv libraries was subjected to four rounds of biopanning against the recombinant SARS-CoV-2 S and RBD proteins (Sino Biological Inc., Beijing, China), fused to mFc or hCκ, as described previously ^35^. Briefly, 3 μg of the recombinant SARS-CoV-2 RBD protein was conjugated to 1.0 × 10^7^ magnetic beads (Dynabeads M-270 epoxy, Invitrogen) and incubated with the scFv phage-display libraries (approximately 10^12^ phages), for 2 h at 37°C. During the first round of biopanning, the beads were washed once with 500 μL 0.05% (v/v) Tween-20 (Sigma-Aldrich, St. Louis, MO, USA) in phosphate-buffered saline (PBST). For the other rounds of biopanning, 1.5 μg recombinant SARS-CoV-2 RBD protein was conjugated to 5.0 × 10^6^ magnetic beads, and the number of washes was increased to three. After each round of biopanning, the bound phages were eluted and rescued, as described previously ^35^.

### Phage ELISA

To select SARS-CoV-2 S reactive clones, phage enzyme-linked immunosorbent assay (ELISA) was performed, using recombinant S and RBD protein-coated microtiter plates, as described previously ^36^. Reactive scFv clones were subjected to Sanger sequencing (Cosmogenetech, Seoul, Republic of Korea), to determine their nucleotide sequences.

### Expression of recombinant proteins

A human, codon-optimized, SARS-CoV-2 RBD (YP_009724390.1, amino acids 306–543) gene was synthesized (Integrated DNA Technologies). Using a synthesized wild-type RBD gene as a template, RBD mutants (V341I, F342L, N354D, N354D/D364Y, V367F, R408I, A435S, W436R, G476S, and V483A) were generated through two-step PCR, using the primers listed in Supplementary Table 8. The genes encoding wild-type or mutant SARS-CoV-2 RBD were cloned into a modified mammalian expression vector, containing the hCκ gene ^35^, and transfected into Expi293F (Invitrogen) cells. The fusion proteins were purified by affinity chromatography, using KappaSelect Columns (GE Healthcare, Chicago, IL, USA), as described previously ^37^. Due to low expression yields, two RBD mutants (N354D/D364Y, W436R) were excluded from further studies.

The genes encoding the selected scFv clones were cloned into a modified mammalian expression vector, containing the hIgG_1_ Fc regions (hFc) or hCκ at the C-terminus ^35,38^, before being transfected and purified by affinity chromatography, as described above.

### ELISA

First, 100 ng of each recombinant SARS-CoV-2 S (Sino Biological Inc.), S1 (Sino Biological Inc.), S2 (Sino Biological Inc.), NP (Sino Biological Inc.), RBD, RBD mutants, SARS-CoV RBD (Sino Biological Inc.), MERS-CoV S (Sino Biological Inc.), RBD (Sino Biological Inc.), S2 (Sino Biological Inc.) proteins were added to microtiter plates (Costar), in coating buffer (0.1 M sodium bicarbonate, pH 8.6). After incubation at 4°C, overnight, and blocking with 3% bovine serum albumin (BSA) in PBS, for 1 h at 37°C, serially diluted plasma (5-fold, 6 dilutions, starting from 1:100) or scFv-hFc (5-fold, 12 dilutions, starting from 1,000 or 500 nM) in blocking buffer was added to individual wells and incubated for 1, h at 37°C. Then, the plates were washed three times with 0.05% PBST. Horseradish peroxidase (HRP)-conjugated rabbit anti-human IgG antibody (Invitrogen) or anti-human Ig kappa light chain antibody (Millipore, Temecula, CA, USA), in blocking buffer (1:5,000), was added into wells and incubated for 1 h at 37°C. After washing three times with PBST, 2,2′-azino-bis-3-ethylbenzothiazoline-6-sulfonic (ThermoFisher Scientific Inc., Waltham, MA, USA) or 3,3′,5,5′-Tetramethylbenzidine liquid substrate system (ThermoFisher Scientific Inc.) was added to the wells. Absorbance was measured at 405 nm or 650 nm, using a microplate spectrophotometer (Multiskan GO; Thermo Scientific).

### Flow cytometry

The recombinant SARS-CoV-2 S protein (200 nM), fused with a polyhistidine tag at the C-terminus (Sino Biological Inc.), was incubated with scFv-hFc fusion proteins at a final concentration of either 200 nM (equimolar) or 600 nM (morlar ratio of 1:3), in 50 μL of 1% (w/v) BSA in PBS, containing 0.02% (w/v) sodium azide (FACS buffer), at 37°C for 1 h. Irrelevant scFv-hFc or scFv-hCκ fusion proteins were used as negative controls. Vero E6 cells (ACE2^+^) were seeded into v-bottom 96-well plates (Corning, Corning, NY, USA), at a density of 1.5 × 10^5^ cells per well. Then, the mixture was added to each well and incubated, at 37°C for 1 h. After washing three times with FACS buffer, FITC-labeled rabbit anti-HIS Ab (Abcam, Cambridge, UK) was incubated, at 37°C for 1 h. Then, the cells were washed three times with FACS buffer, resuspended in 150 μL of PBS, and subjected to analysis by flow cytometry, using a FACS Canto II instrument (BD Bioscience, San Jose, CA, USA). For each sample, 10,000 cells were assessed.

### Microneutralization assay

The virus (BetaCoV/Korea/SNU01/2020, accession number MT039890) was isolated at the Seoul National University Hospital and propagated in Vero cells (ATCC CCL-81), using Dulbecco’s Modified Eagle’s Medium (DMEM, Welgene, Gyeongsan, Republic of Korea) supplemented with 2% fetal bovine serum (Gibco) ^39^. The cells were grown in T-25 flasks, (ThermoFisher Scientific Inc.), inoculated with SARS-CoV-2, and incubated at 37°C, in a 5% CO_2_ environment. Then, 3 days after inoculation, the viruses were harvested and stored at −80°C. The virus titer was determined via a TCID_50_ assay ^40^.

Vero cells were seeded in T-25 flasks and grown for 24 h, at 37°C, in a 5% CO_2_ environment, to ensure 80% confluency on the day of inoculation. The recombinant scFv-hCκ fusion proteins (0.5, 5, or 50 μg/mL) were mixed with 2,500 TCID_50_ of SARS-CoV-2, and the mixture was incubated for 2 h, at 37°C. Then, the mixture (1 mL) was added to the Vero cells and incubated for 1 h, at 37°C, in a 5% CO_2_ environment. After incubation for 1 h, 6 mL of complete media was added to the flasks and incubated, at 37°C, in a 5% CO_2_ environment. After 0, 24, 48, and 72 h of infection, the culture supernatant was collected, to measure the virus titers. RNA was extracted, using the MagNA Pure 96 DNA and Viral NA small volume kit (Roche, Germany), according to the manufacturer’s instructions. Viral RNA was detected using the PowerChek 2019-nCoV Real-time PCR Kit (Kogene Biotech, Seoul, Republic of Korea), for the amplification of the E gene, and quantified according to a standard curve, which was constructed using *in vitro* transcribed RNA, provided by the European Virus Archive (https://www.european-virus-archive.com).

## Data and materials availability

Raw sequencing data will be submitted shortly. Computer codes and processed data will be deposited on Github. All other data that supporting the findings of this study are available from the corresponding author on reasonable request.

## Acknowledgments

The authors thank Su Jin Choi for technical support. This research was funded by the National Research Foundation of Korea [NRF-2016M3A9B6918973] and the Ministry of Science and ICT(MSIT) of the Republic of Korea and the National Research Foundation of Korea [NRF-2020R1A3B3079653]. This research was supported by the Global Research Development Center Program, through the NRF, funded by the MSIT [2015K1A4A3047345]. This work was supported by the Brain Korea 21 Plus Project in 2020.

## Author contributions

SangIl K. designed and conducted the experiments, performed analysis, interpreted experimental results, and wrote the paper. J.N. performed the bioinformatic analysis, visualized and interpreted results, and wrote the paper. Sujeong K. conducted experiments, performed analyses, and interpreted experimental results. Y.C., D.Y., and M.S. conducted the experiments. Y.L. and H.L. performed the bioinformatic analysis. J.J., C.K., K.S., P.C., H.K., E.K., and N.K. contributed to patient recruitment. W.P. conceived the study, designed and conducted experiments, and interpreted experimental results. M.O. conceived the study. S.K. conceived the study and designed and supervised the bioinformatics analysis. J.C. conceived the study, designed and supervised the experiments, interpreted all results, and wrote the paper.

---

**Supplementary Fig 1.**
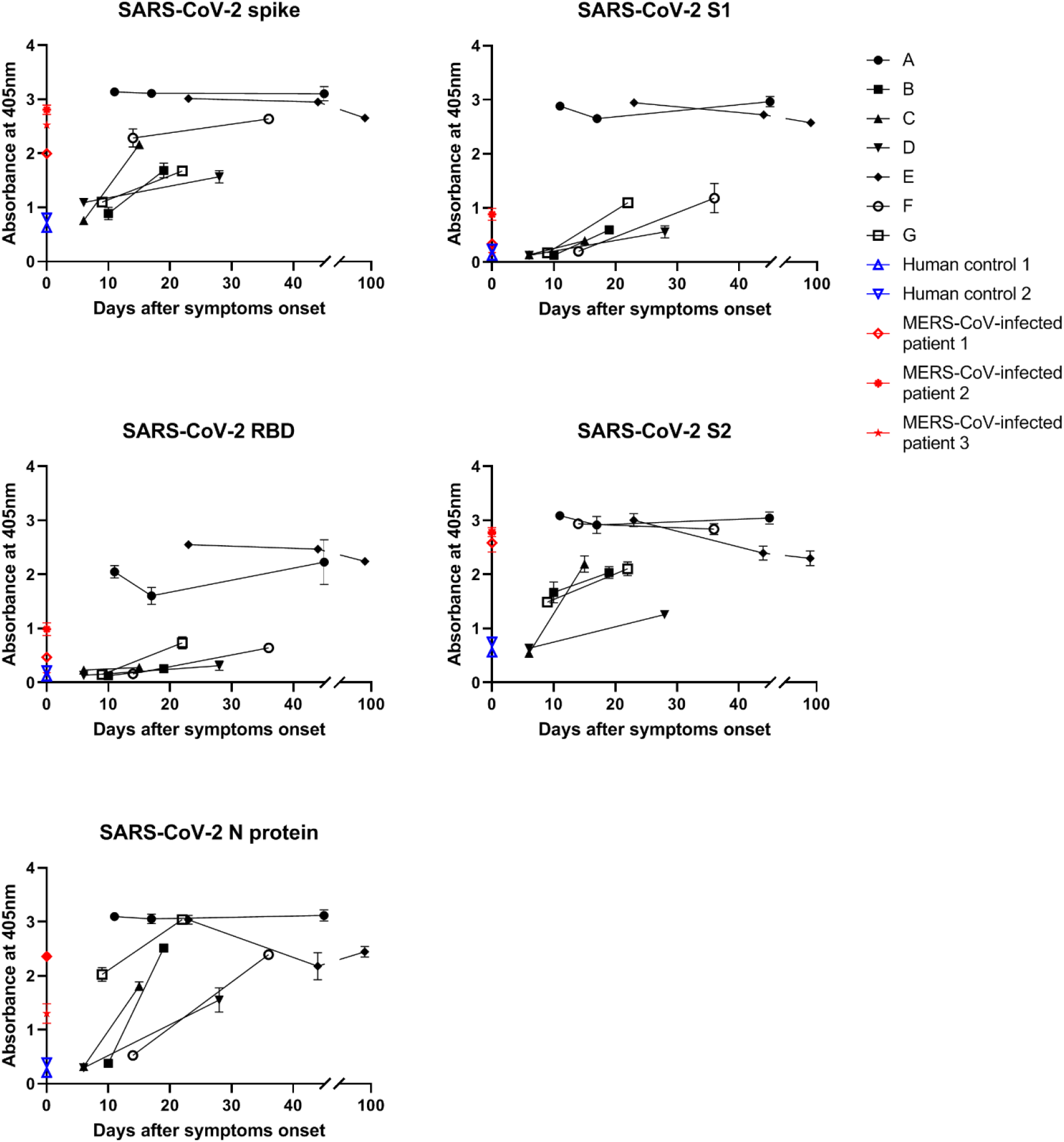
Titrations of serum IgG in ELISA. Plasma samples from seven SARS-CoV-2 patients were diluted (1:100) and added to plates coated with recombinant SARS-CoV-2 spike, S1, S2, or N proteins, fused to HIS tag, or RBD protein, fused to human Cκ domain. The amount of bound IgG was determined using anti-human IgG (Fc-specific) antibody. ABTS was used as the substrate. All experiments were performed in duplicate, and the data are presented as the mean ± SD.

**Supplementary Fig 2.**
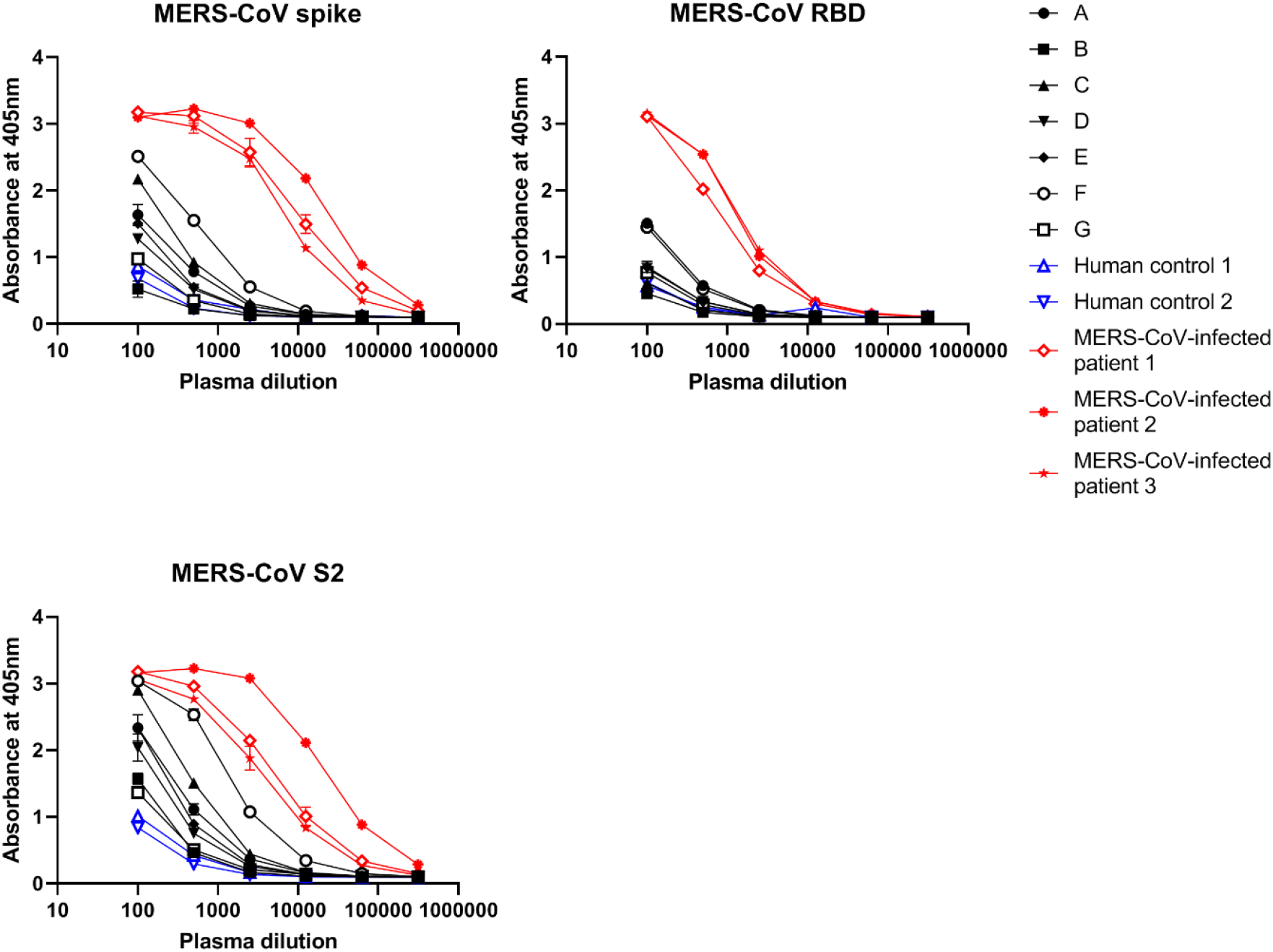
Titrations of serum IgG in ELISA. Plasma samples of seven SARS-CoV-2 patients were serially diluted and added to plates coated with recombinant MERS-CoV spike, RBD, and S2 proteins, fused to HIS. The amount of bound IgG was determined using anti-human IgG (Fc-specific) antibody. ABTS was used as the substrate. All experiments were performed in duplicate, and the data are presented as the mean ± SD.

**Supplementary Fig 3.**
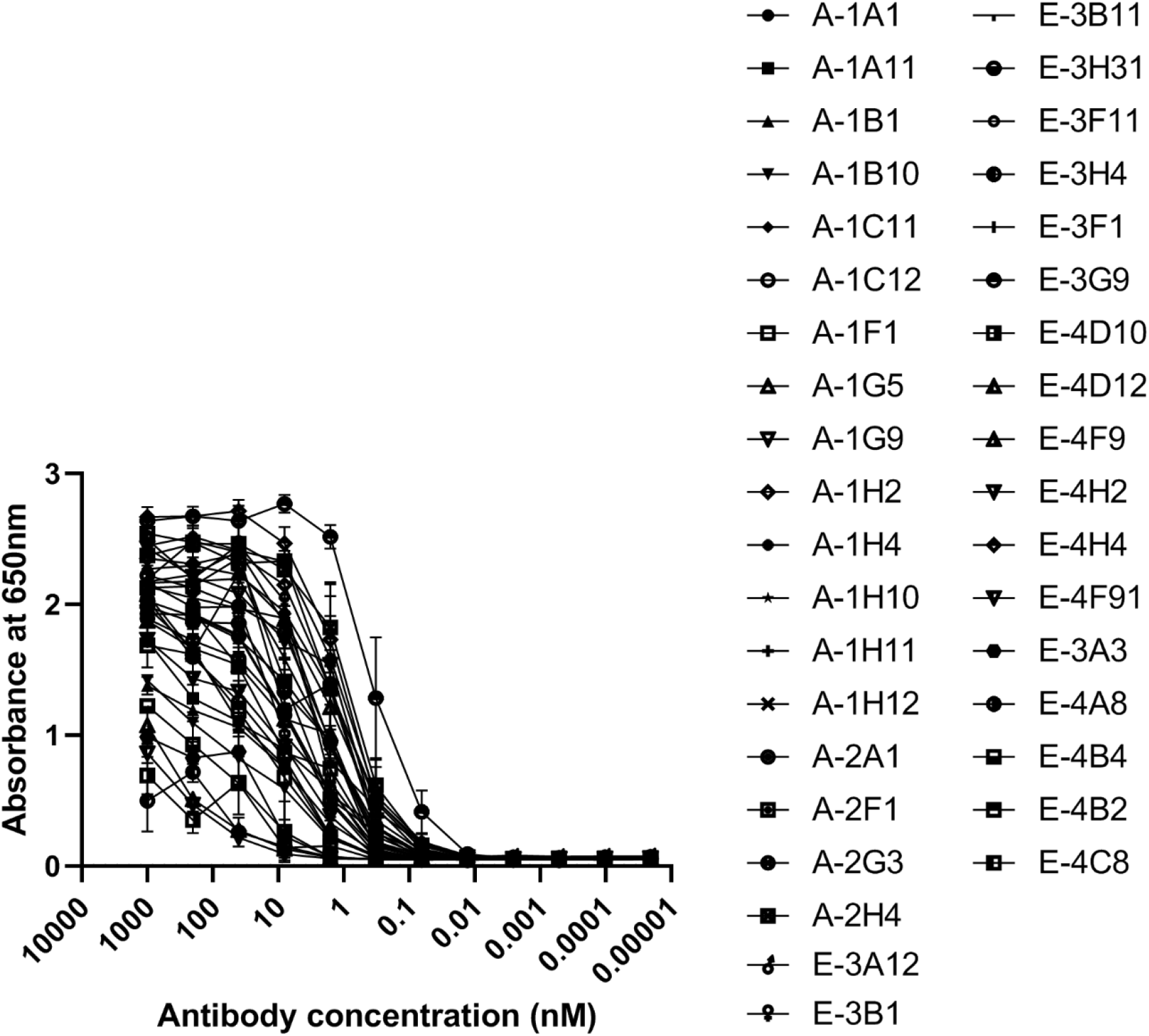
Reactivity of anti-SARS-CoV-2 scFv antibodies against recombinant SARS-CoV-2 RBD. Recombinant SARS-CoV-2 RBD-coated microtiter plates were incubated with varying concentrations of scFv-hCκ fusion proteins. HRP-conjugated anti-human Ig kappa light chain antibody was used as the probe, and TMB was used as the substrate. All experiments were performed in duplicate, and the data are presented as the mean ± SD.

**Supplementary Fig 4.**
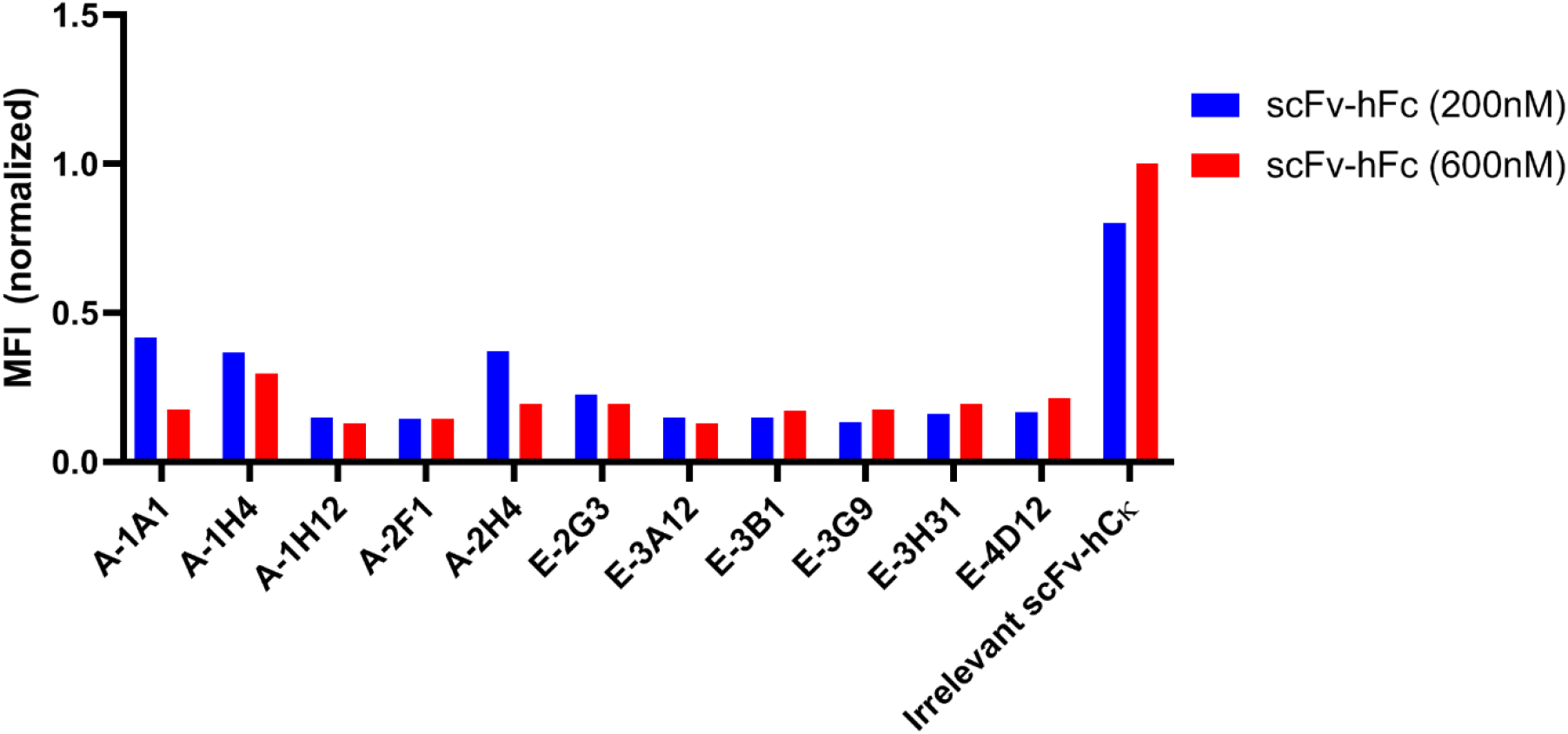
Inhibition of recombinant SARS-CoV-2 S glycoprotein binding to ACE2-expressing cells, by flow cytometry. The recombinant scFv-hFc fusion proteins (200 nM or 600 nM) were mixed and incubated with recombinant SARS-CoV-2 S glycoprotein (200 nM) fused with a HIS tag at the C-terminus. After incubation with Vero E6 (ACE2^+^) cells, the relative amount of bound, recombinant SARS-CoV-2 S glycoprotein was measured using a FITC-conjugated anti-HIS antibody. For each sample, 10,000 cells were monitored.

**Supplementary Fig 5.**
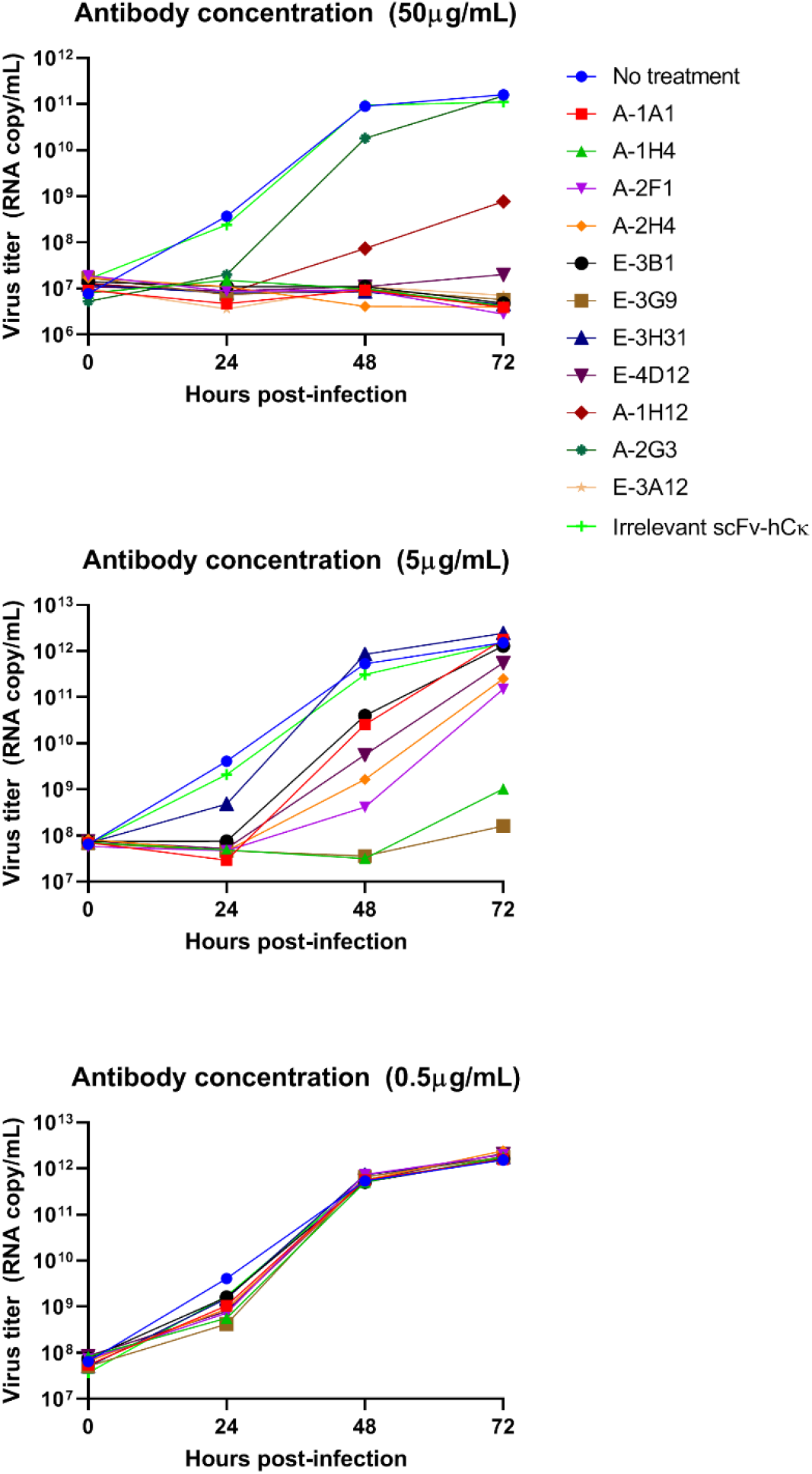
Neutralization of SARS-COV-2 in an *in vitro* experiment. The recombinant scFv-hCκ fusion proteins were mixed with 2,500 TCID_50_ of SARS-CoV-2 (BetaCoV/Korea/SNU01/2020, accession number MT039890), and the mixture was added to the Vero cells. After 0, 24, 48, and 72 h of infection, the culture supernatant was collected to measure the viral titers.

**Supplementary Fig 6.**
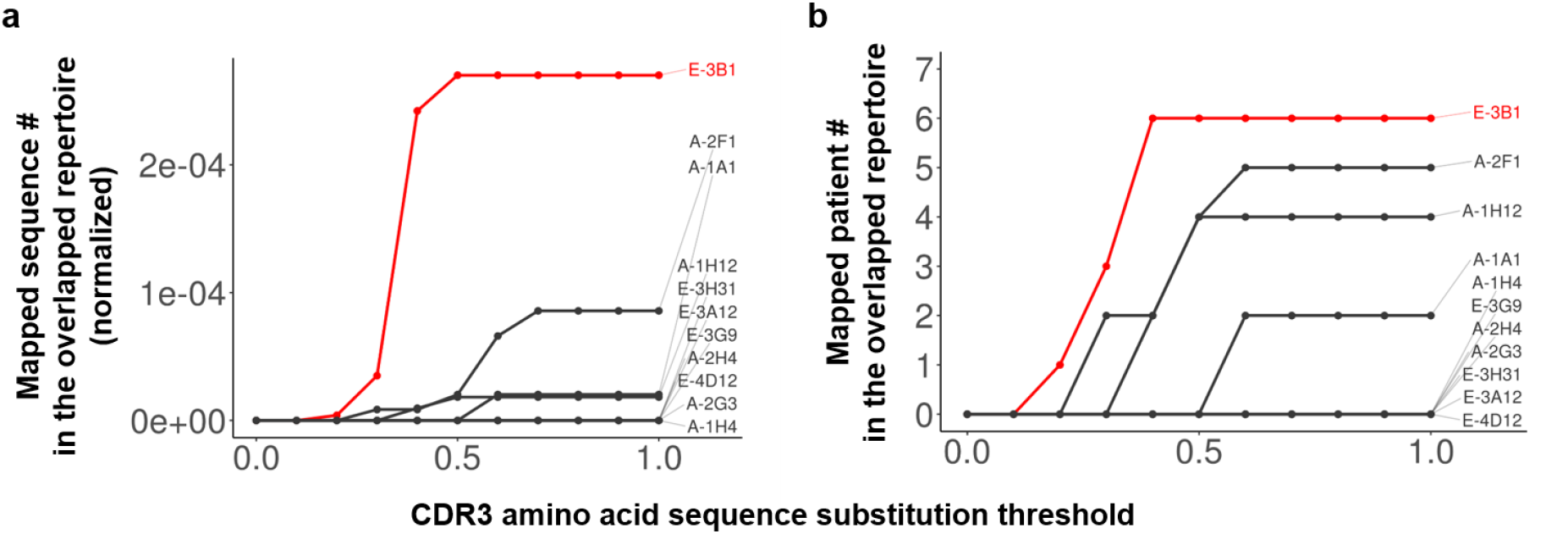
Mapping of the 11 nAbs to the overlapping IGH repertoire. **a,** The number class-switched IGH sequences in the overlapping repertoire, mapped to nAbs. The allowed number of CDR3 amino acid sequence substitutions during the mapping process is represented on the x-axis of the plot, after normalizing against the sequence length. The number of mapped sequences was normalized against the total number of IGH sequences in each patient, and their sum is represented in the y-axis of the plot. **b,** The number of patients expressing the overlapping class-switched IGH sequences, which were mapped to the nAbs. The x-axis is the same as described for **a** and the y-axis indicates the number of patients.

**Supplementary Fig 7.**
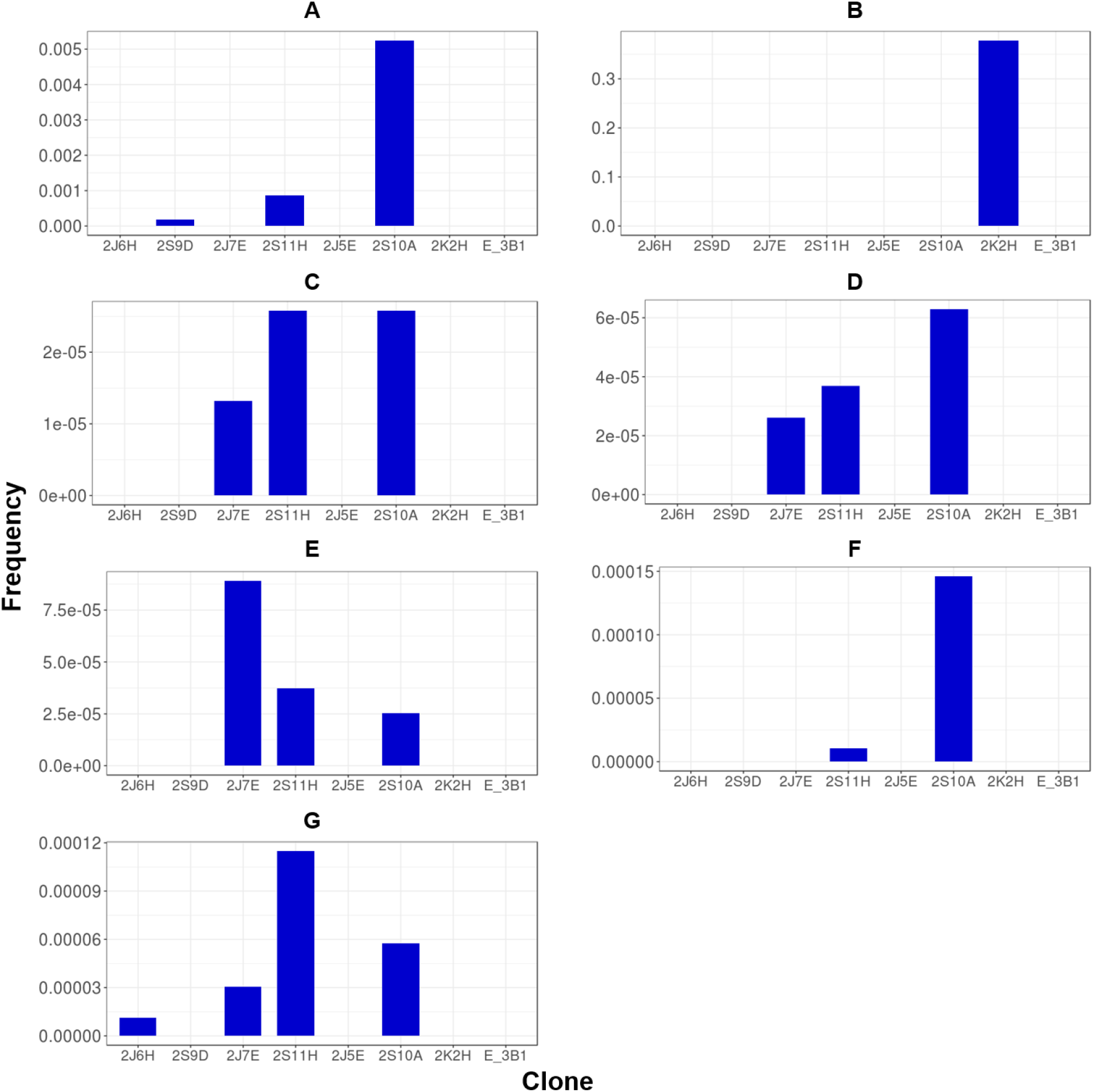
Existence of V_L_ that can be paired with the stereotypic V_H_. V_L_ was mapped according to identical VJ gene usage and perfectly matched LCDR3 sequences at the amino acid level, which were identified in the IGL repertoires of all seven patients (A–G). The frequency values of the mapped sequences in the repertoires of all sampling points were summed. Patient identification can be found above each bar graph.

**Supplementary Fig 8.**
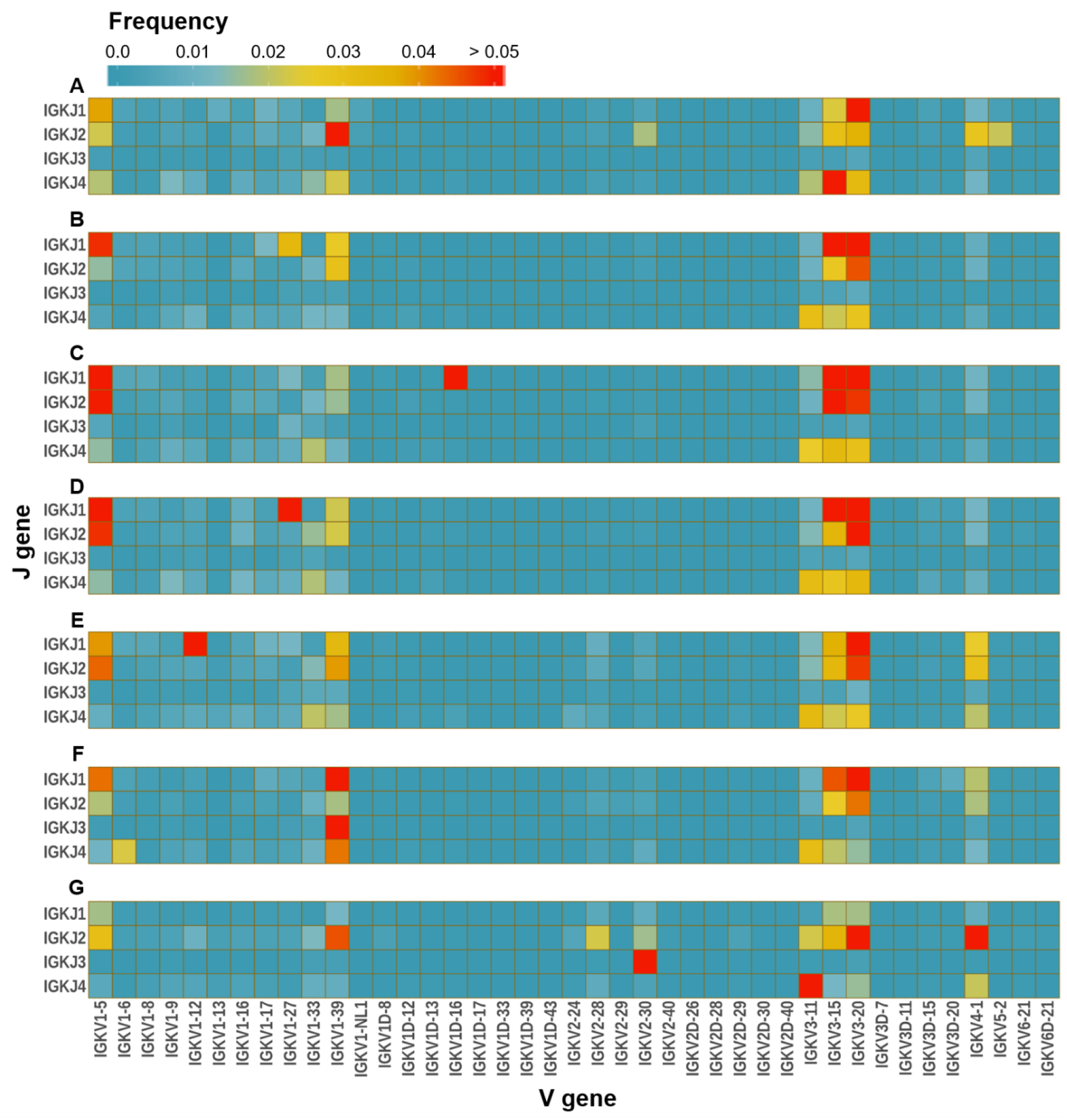
VJ gene usage among the IG kappa light chain repertoire of patients. The frequency values of all sampling points were averaged and represented for each patient. Patient identification can be found at the top left corner of each heatmap.

**Supplementary Fig 9.**
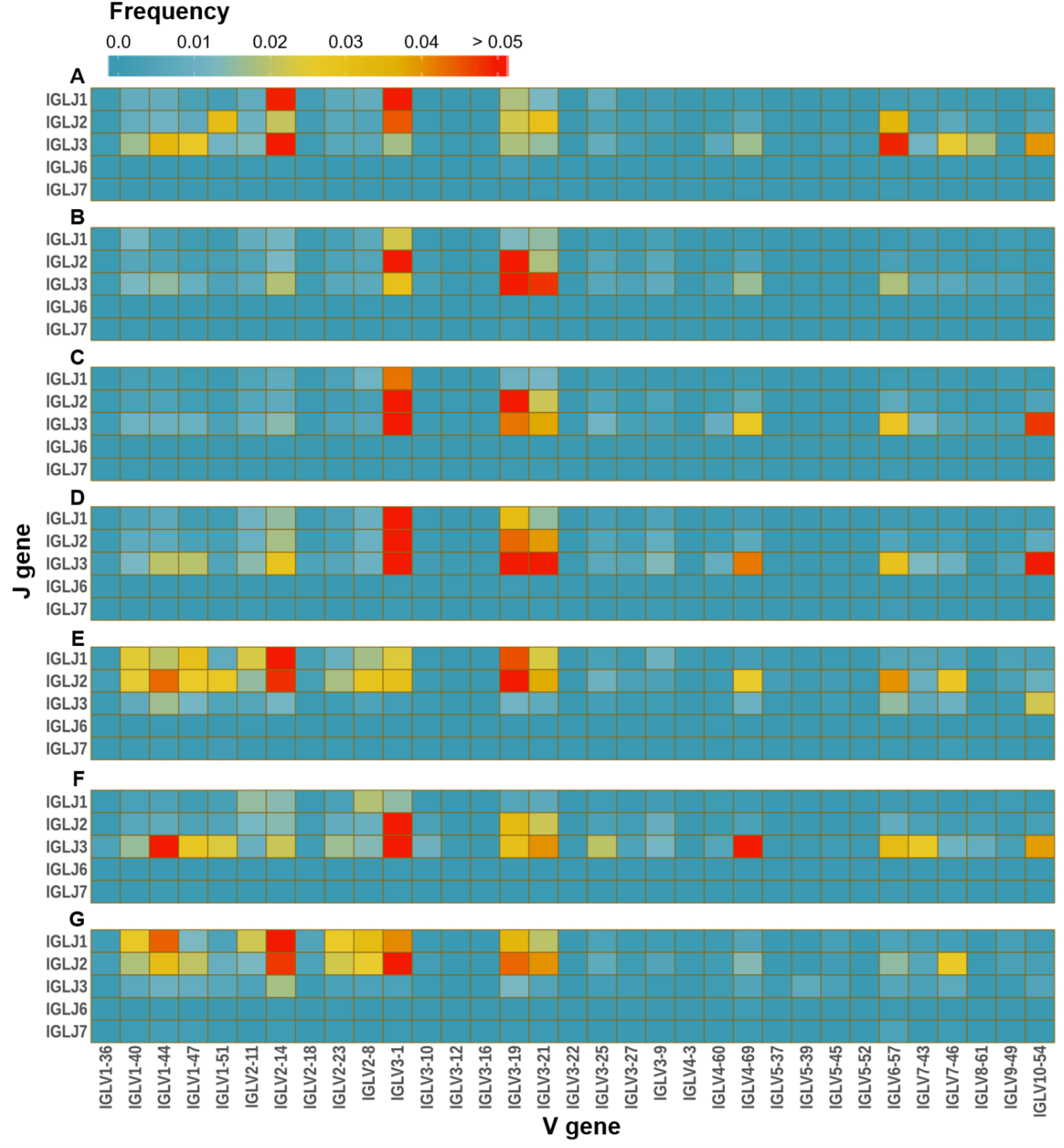
VJ gene usage among the IG lambda light chain repertoire of patients. The frequency values of all sampling points were averaged and are represented for each patient. Patient identification can be found at the left top corner of each heatmap.

**Supplementary Fig 10.**
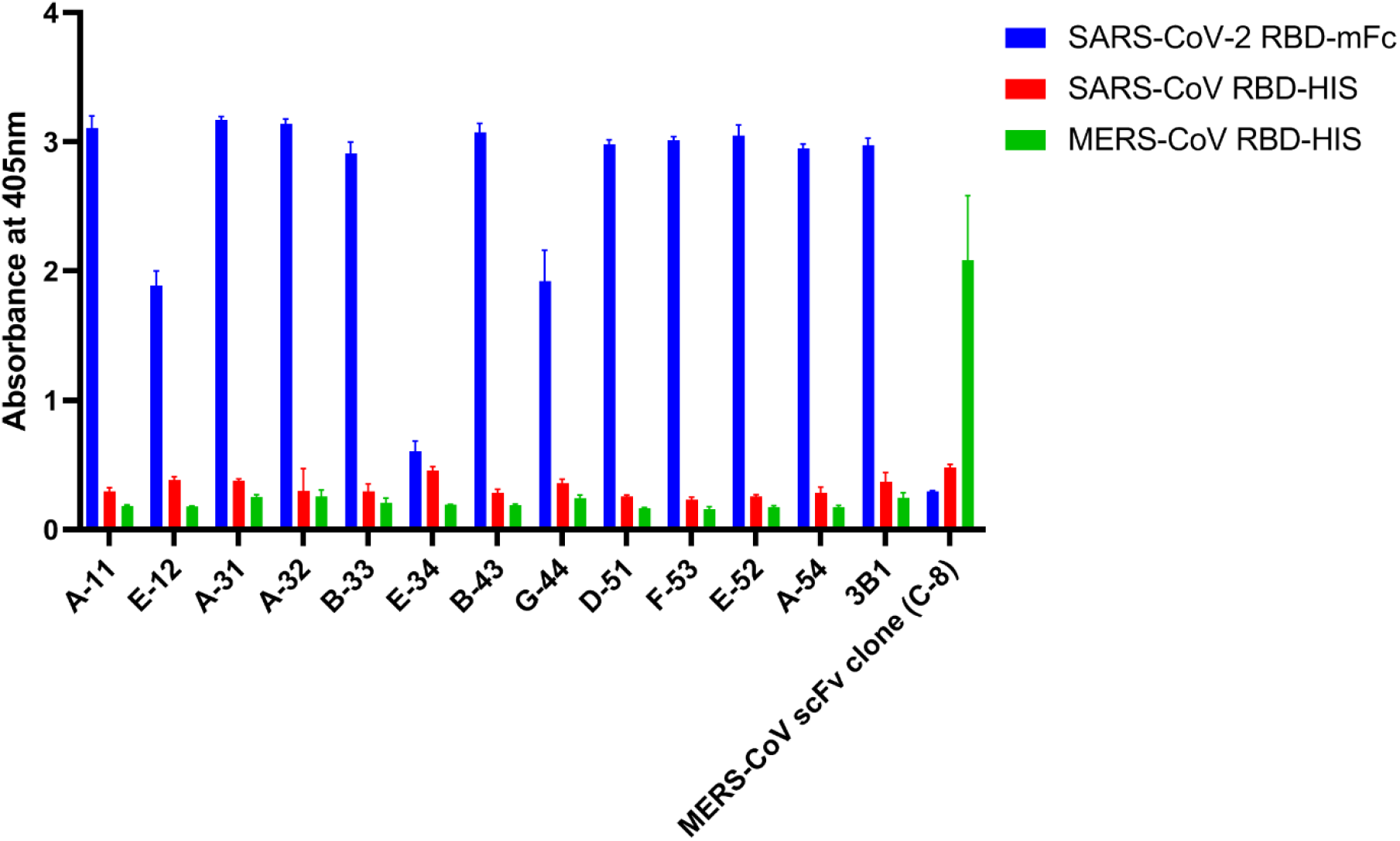
Reactivity of phage-displayed scFv clones in phage ELISA. Recombinant SARS-CoV-2, SARS-CoV, or MERS-CoV RBD protein-coated microtiter plates were incubated with phage clones. HRP-conjugated anti-M13 antibody was used as the probe, and ABTS was used as the substrate. All experiments were performed in quadruplicate, and the data are presented as the mean ± SD.

**Supplementary Fig 11.**
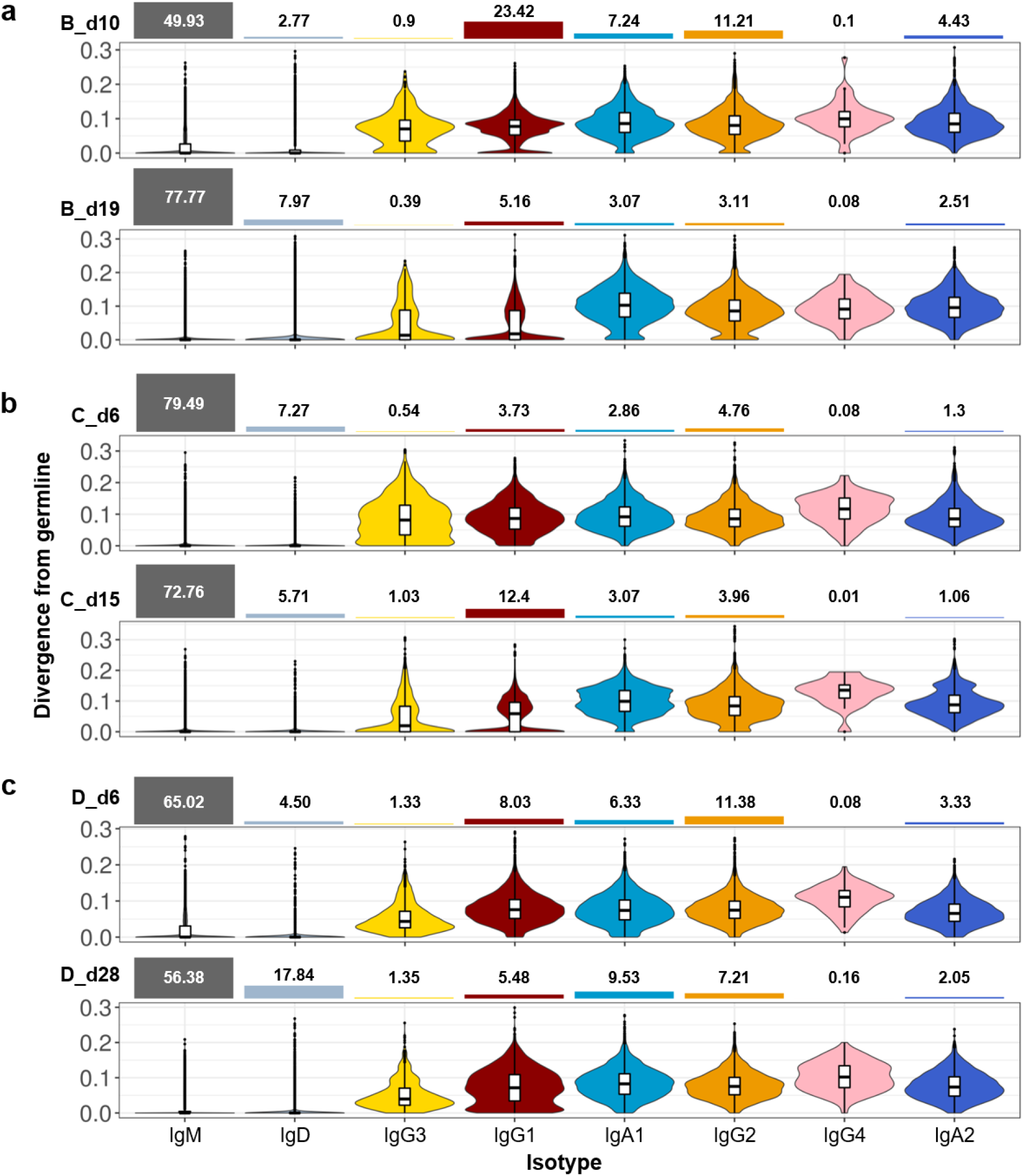
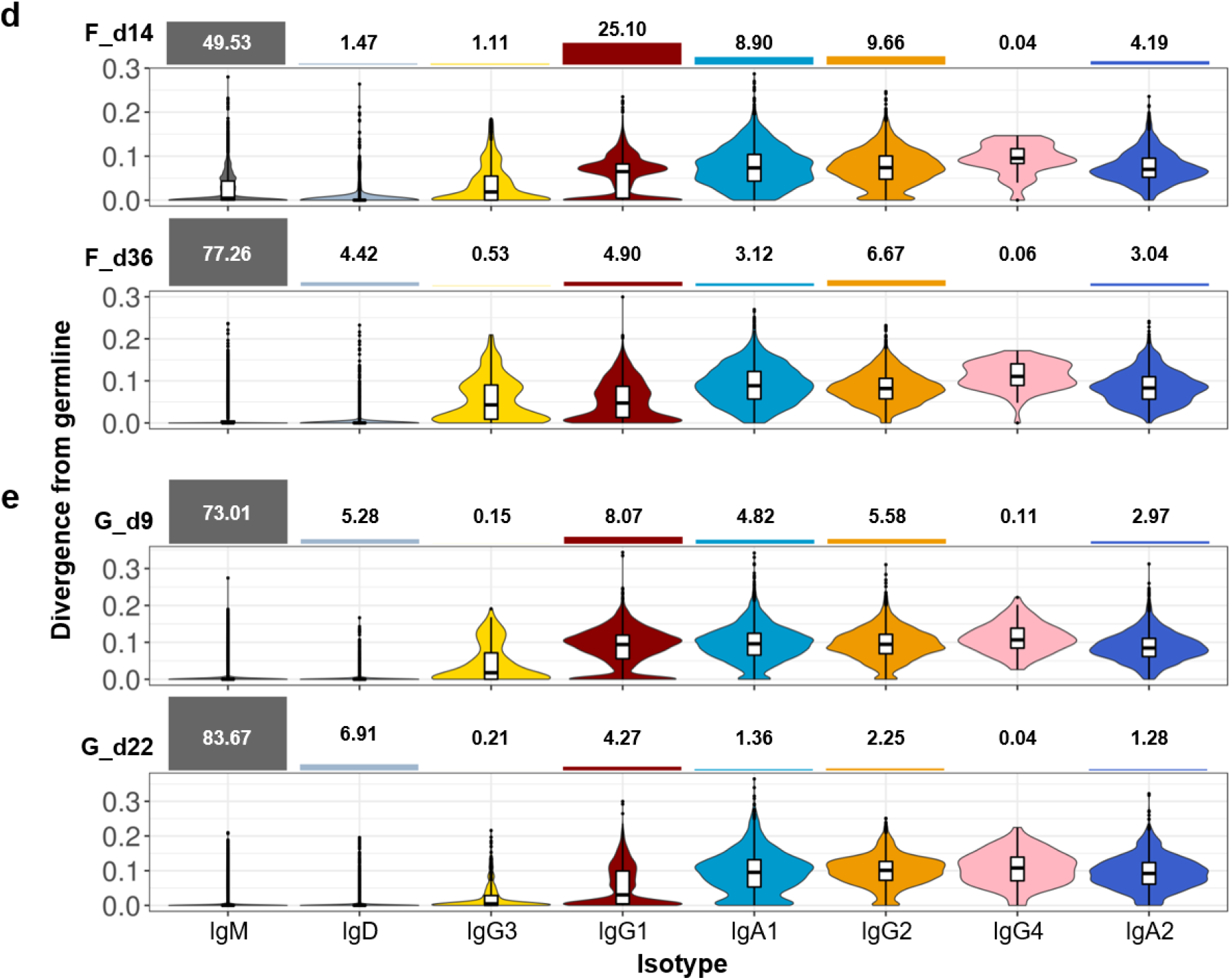
Deep profiling of the IGH repertoire of Patients B, C, D, F, and G. **(a-e),** IGH repertoires of **a.** Patient B, **b.** Patient C, **c.** Patient D, **d.** Patient F, and **e.** Patient G were examined according to divergence from the germline and the isotype composition of the sequences. Values of divergence from the germline were calculated separately, for each isotype, and are presented as violin plots, class-switching event order. The bar graphs above the violin plots represent the proportions of each isotype.

**Supplementary Fig 12.**
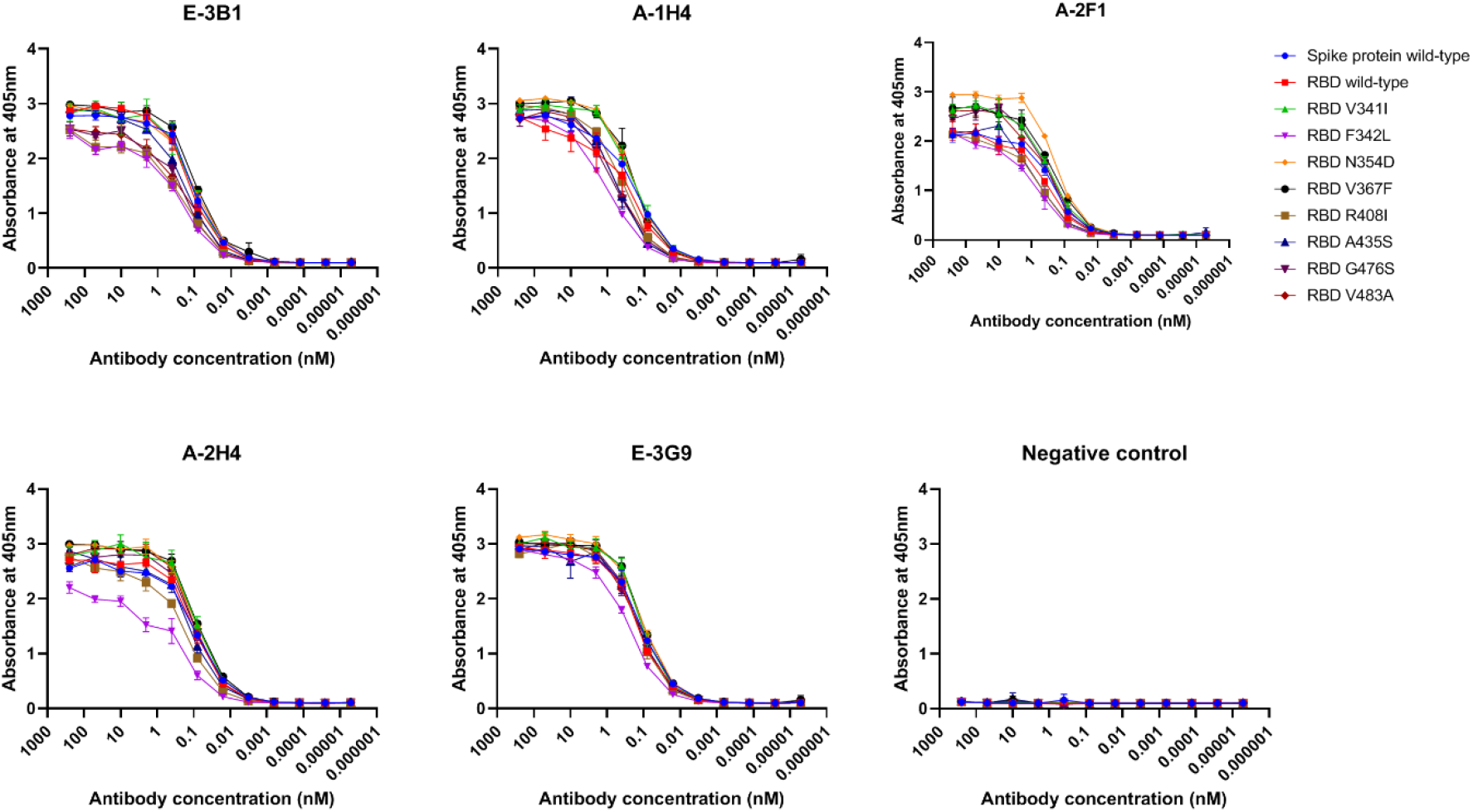
Reactivity of nAbs against recombinant SARS-CoV-2 RBD mutants. Recombinant wild-type or mutant (V341I, F342L, N354D, V367F, R408I, A435S, G476S, and V483A) SARS-CoV-2 RBD protein-coated microtiter plates were incubated with varying concentrations of scFv-hFc fusion proteins. HRP-conjugated anti-human IgG antibody was used as the probe, and ABTS was used as the substrate. All experiments were performed in triplicate, and the data are presented as the mean ± SD.

**Supplementary Fig 13.**
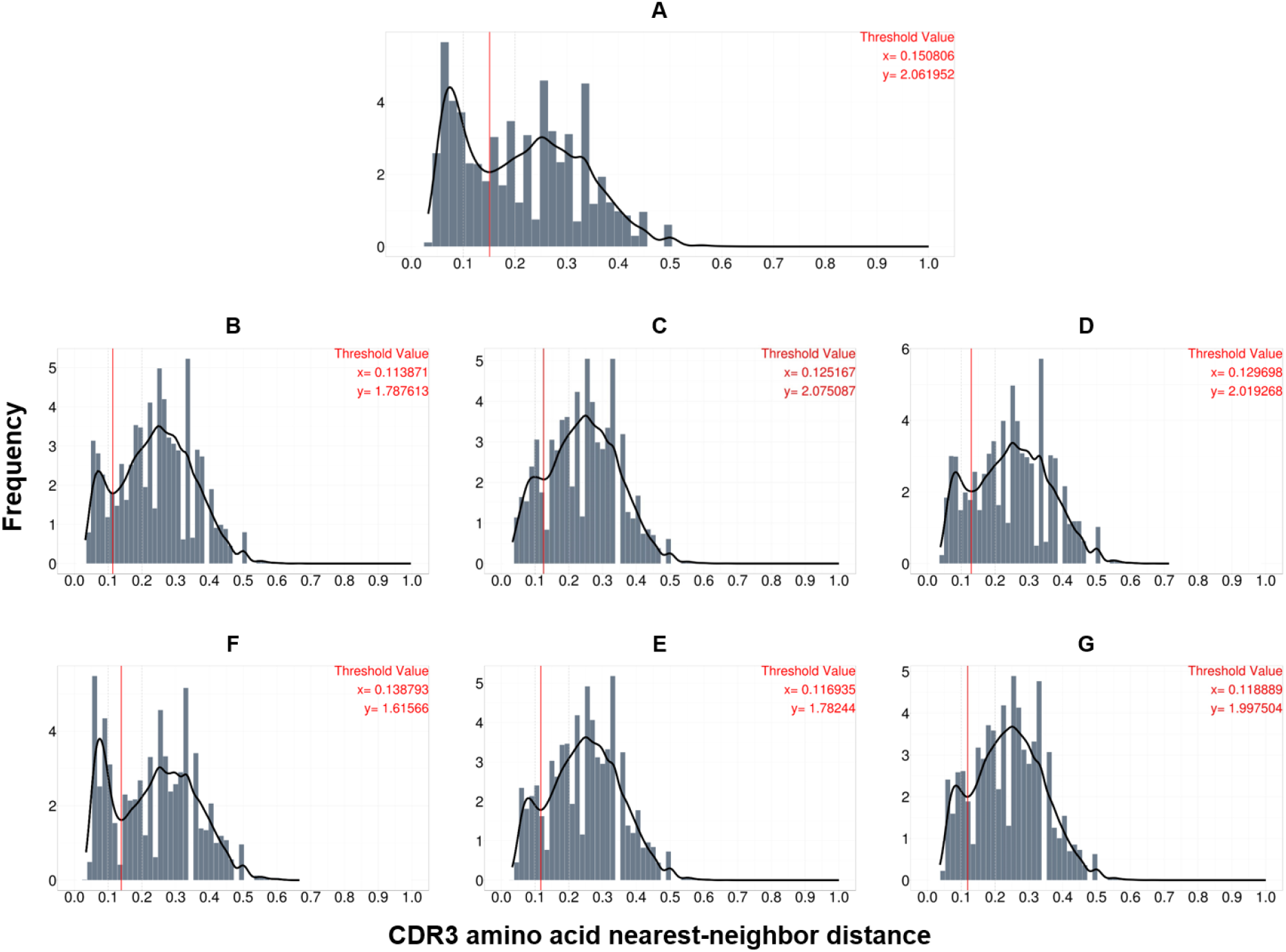
The nearest-neighbor distance histogram for HCDR3 amino acid sequences in the IGH repertoires of patients. The frequency values of the histograms were approximated by the binned kernel estimation method, in the Gaussian kernel setting (black line). The threshold value for each patient was set as the x value of the points with a minimum frequency value between two peaks of the bimodal distribution (red vertical line). The x and y values of the threshold-setting point are indicated in the upper right corner of each histogram.

**Supplementary Fig 14.**
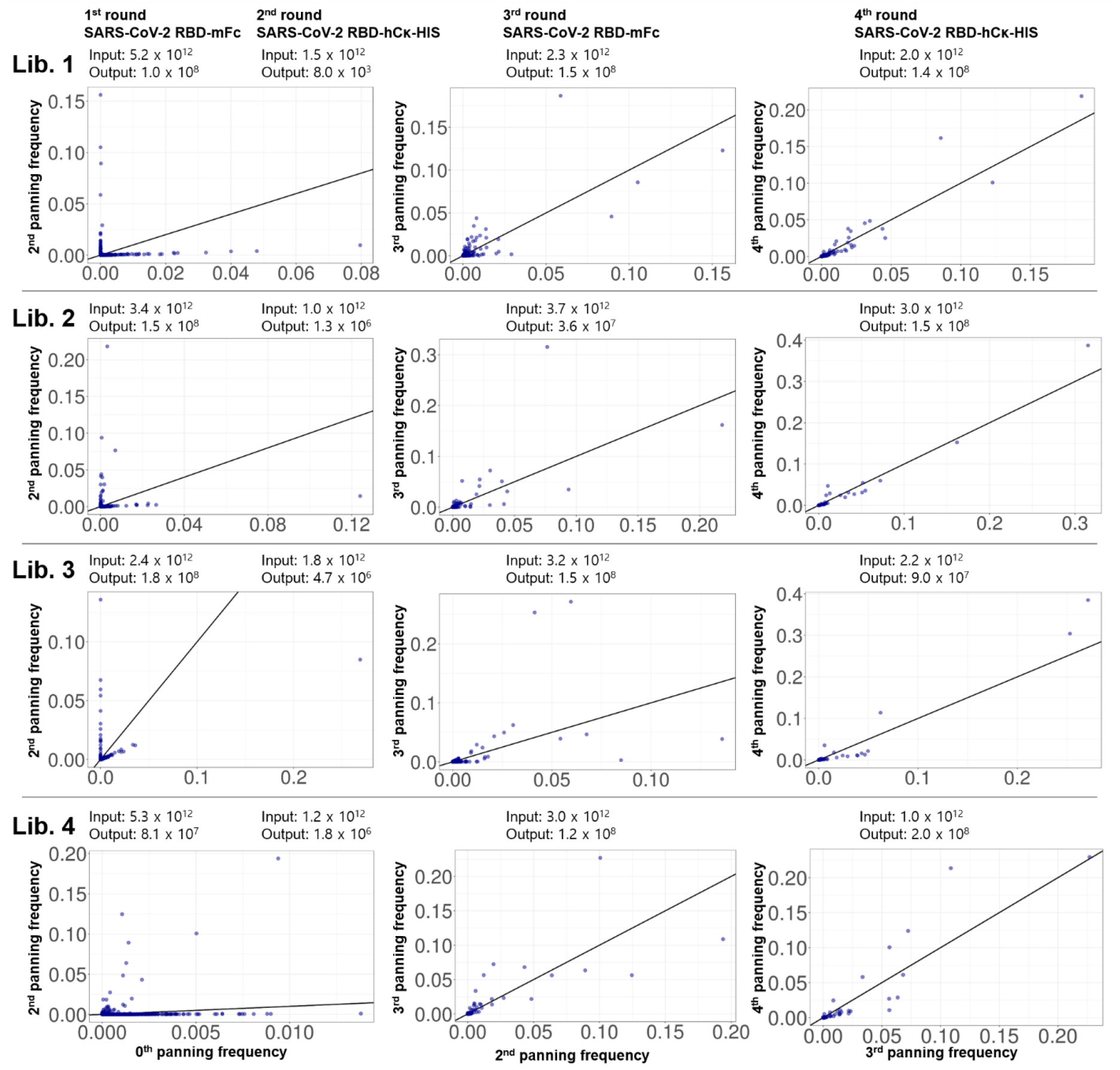
Frequency scatter plots for the NGS data of the four libraries, after each round of biopanning. The x- and y-axes represent the frequency values for the NGS data in each biopanning round. The line on the scatter plots indicates the identity line (y = x). Input and output virus titer values are also presented, above the matched scatter plots.

**Supplementary Fig 15.**
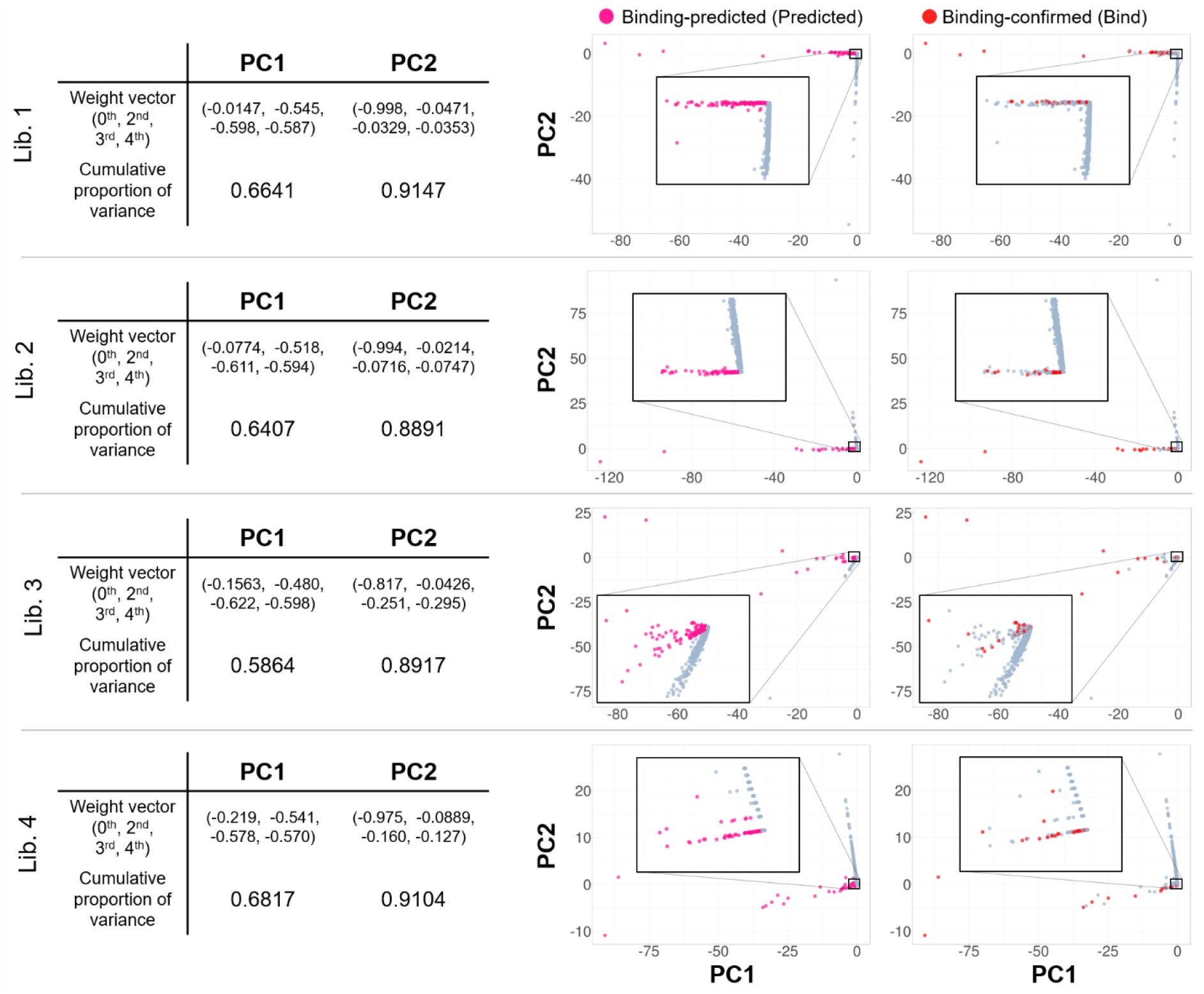
The results of principal component analysis, applied to the NGS data of four libraries, after each round of biopanning. Information regarding the PC weight vectors and the cumulative proportion of variance explained by the PCs are listed on the left side of the plots. PCA plots for PC1 and PC2 on shown on the right side of the plots. The binding-predicted clones were defined based on the value of PC1 and the ratio between PC1 and PC2, by setting a constant threshold value for each. The population of clones defined as predicted clones is marked in pink. The clones known bind to SARS-CoV-2 RBD are marked in red.

**Supplementary Table 1.** Demographic and clinical characteristics.

| Patient no. | A | B | C | D | E | F | G |
| --- | --- | --- | --- | --- | --- | --- | --- |
| Age | 55 | 55 | 53 | 24 | 48 | 40 | 59 |
| Sex | Male | Male | Female | Male | Male | Female | Female |
| Race | Korean | Korean | Korean | Korean | Chinese | Chinese | Korean |
| BMI (kg/m<sup>2</sup>) | 31.35 | 24.09 | 23.10 | 21.51 | 27.02 | 22.15 | 18.00 |
| Underlying diseases | - | DM, HTN, DL | - | - | - | - | DM, DL |
| Highest temperature (°C) | 39.7 | 38.4 | 38.0 | 37.8 | 37.8 | 37.8 | 37.0 |
| Symptoms | Dyspnea, myalgia, diarrhea | Sputum, myalgia | Sputum, myalgia | Myalgia | Cough, myalgia, diarrhea | Cough, sputum, myalgia, diarrhea | - |
| Pneumonic infiltrates | Extensive | Limited | Limited | Limited | Extensive | Limited | Limited |
| Oxygen therapy | Yes | No | No | No | No | No | No |
| Ventilator | No | No | No | No | No | No | No |
| Antiviral Treatment | Lopinavir/ritonavir | - | - | - | Lopinavir/ritonavir | Lopinavir/ritonavir | - |
| Antibiotic Treatment | Levofloxacin | - | - | - | - | - | - |
| Blood samples collected after symptoms onset (Days) | 11, 17, 45 | 10, 19 | 6, 15 | 6, 28 | 23, 44, 99 | 14, 36 | 9, 22 |

BMI, body mass index; DM, diabetes mellitus; HTN, hypertension; DL, dyslipidemia

**Supplementary Table 2.**
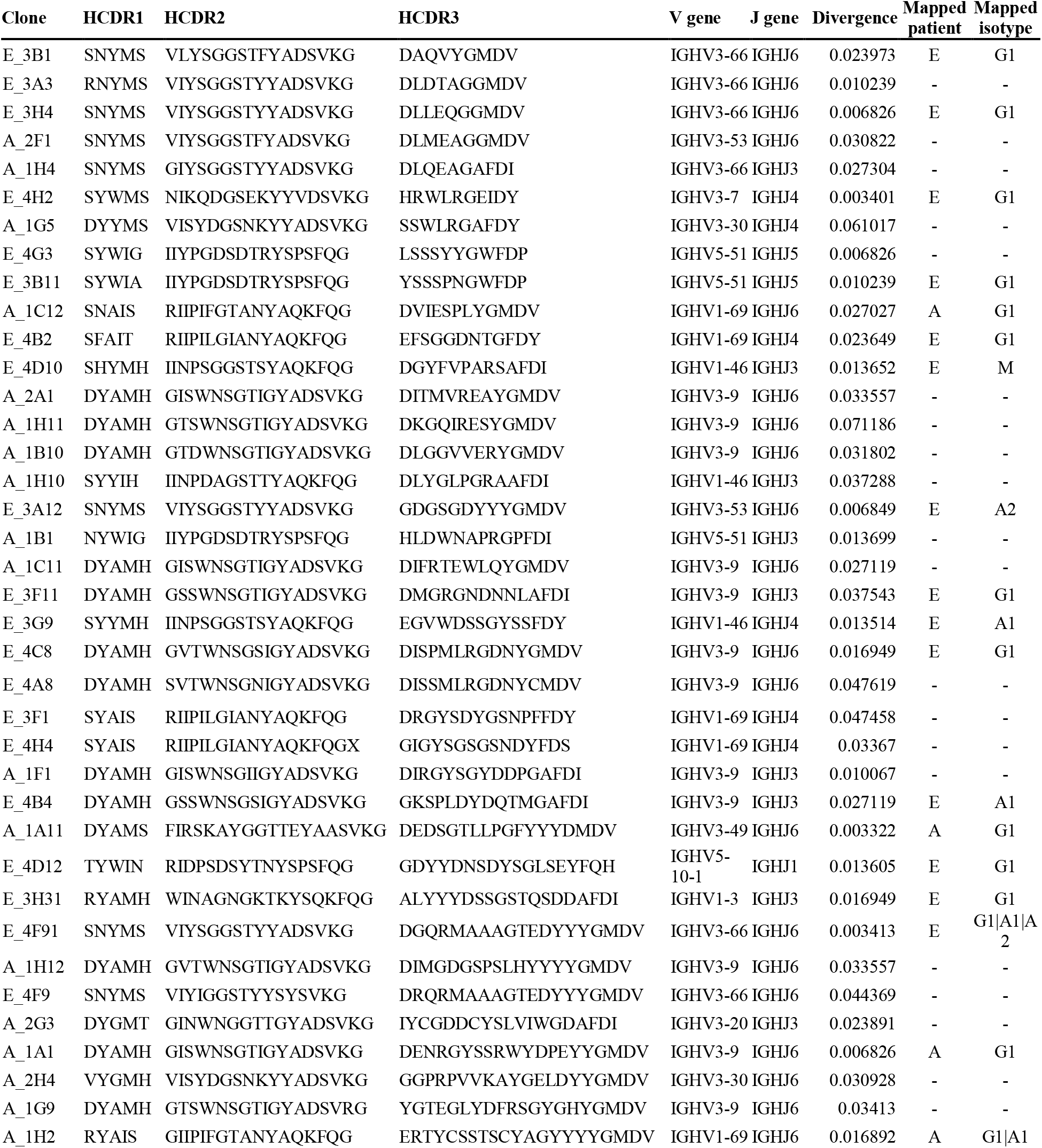
SARS-CoV-2 RBD-reactive scFv clones.

**Supplementary Table 3.**
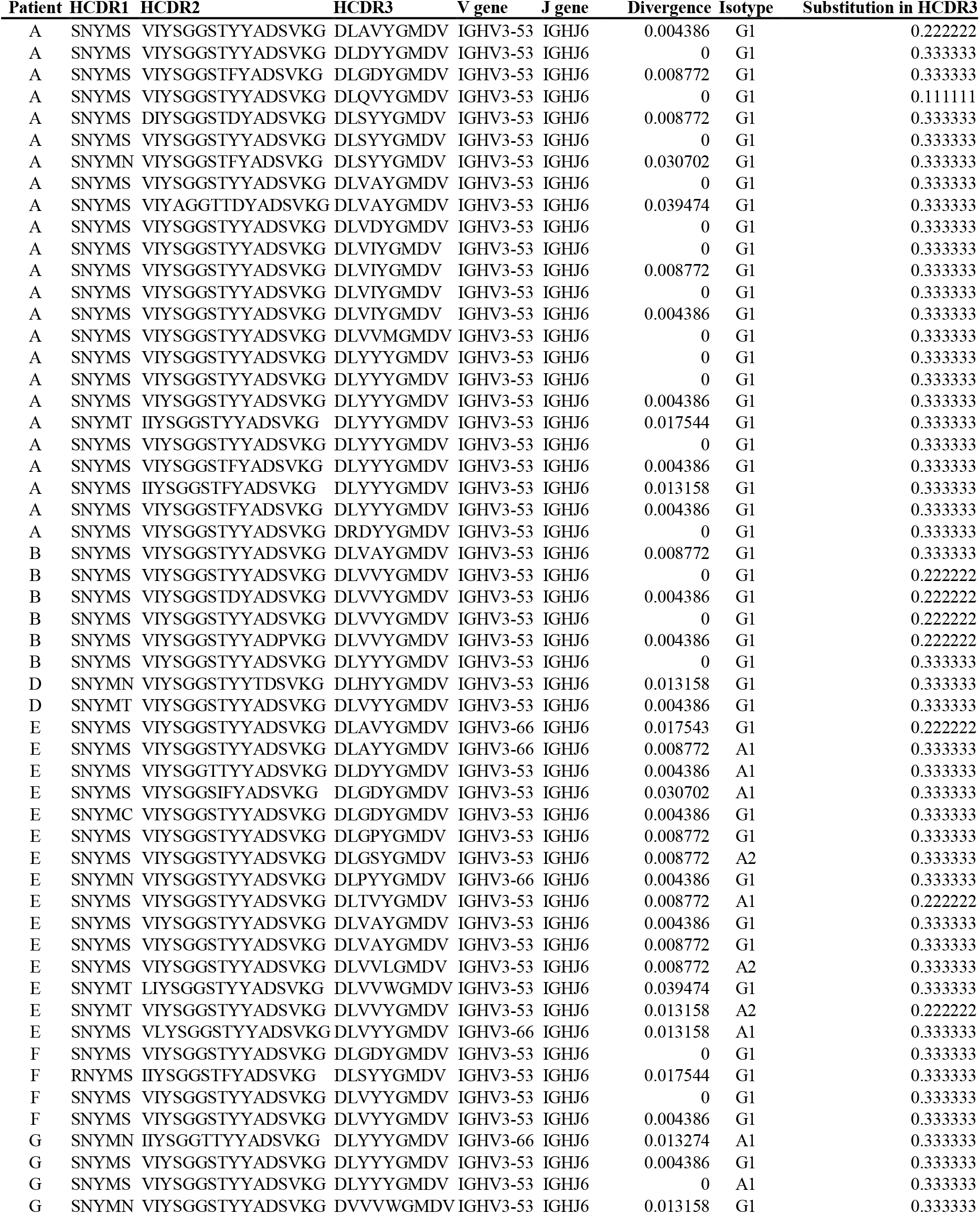
Class-switched IGH clonotypes homologous to E-3B1.

**Supplementary Table 4.** Human monoclonal antibodies reactive against MERS-CoV RBD.

| Clone | V gene | J gene | Divergence |
| --- | --- | --- | --- |
| 4 | IGHV3-23 | IGHJ4 | 0.075862 |
| 13 | IGHV3-23 | IGHJ4 | 0.061224 |
| 28 | IGHV3-30 | IGHJ6 | 0.013559 |
| 34 | IGHV3-21 | IGHJ3 | 0.057627 |
| 36 | IGHV3-23 | IGHJ4 | 0.064626 |
| 38 | IGHV3-23 | IGHJ4 | 0.088136 |
| 42 | IGHV3-21 | IGHJ4 | 0.061433 |
| 103 | IGHV3-23 | IGHJ4 | 0.050847 |
| 119 | IGHV3-9 | IGHJ6 | 0.087838 |
| 180 | IGHV6-1 | IGHJ4 | 0.009836 |
| 113 | IGHV3-30 | IGHJ4 | 0.00339 |
| 121 | IGHV1-69 | IGHJ4 | 0.128028 |
| 6 | IGHV4-39 | IGHJ5 | 0.016892 |
| 25 | IGHV4-39 | IGHJ5 | 0.016892 |
| 10-1 | IGHV1-69 | IGHJ5 | 0.006757 |
| 20-1 | IGHV1-69 | IGHJ5 | 0.006757 |
| 38-1 | IGHV1-69 | IGHJ5 | 0.006757 |
| 39 | IGHV1-69 | IGHJ5 | 0.010135 |
| 40 | IGHV1-69 | IGHJ5 | 0.006757 |
| 11 | IGHV4-39 | IGHJ4 | 0.020067 |
| 26 | IGHV4-39 | IGHJ4 | 0.036789 |
| 21 | IGHV1-69 | IGHJ5 | 0.003378 |
| 17 | IGHV1-69 | IGHJ3 | 0.010135 |
| 30 | IGHV1-69 | IGHJ3 | 0.020339 |
| 33 | IGHV1-69 | IGHJ3 | 0.016949 |
| 41 | IGHV1-69 | IGHJ3 | 0.013514 |
| 46 | IGHV4-39 | IGHJ4 | 0.016722 |
| 47 | IGHV4-39 | IGHJ4 | 0.016722 |
| 48 | IGHV4-39 | IGHJ4 | 0.020067 |
| 7 | IGHV3-21 | IGHJ6 | 0.010274 |
| 9 | IGHV1-69 | IGHJ4 | 0.017065 |
| 31 | IGHV1-69 | IGHJ5 | 0.020339 |
| 35 | IGHV3-21 | IGHJ6 | 0.006849 |
| 42-1 | IGHV4-39 | IGHJ5 | 0.020067 |
| 10 | IGHV4-39 | IGHJ5 | 0.016722 |
| 15 | IGHV1-69 | IGHJ4 | 0.003413 |
| 20 | IGHV1-69 | IGHJ4 | 0.020619 |

**Supplementary Table 5.** Statistics for the pre-processing of the IGH NGS data.

| **Sample** | **Raw read** | **UMI-processed read**<br>**(functional filtering**<br>**performed)** | **Unique consensus**<br>**sequence #** | **Sampled UMI-**<br>**processed read** | **Sampled unique**<br>**consensus**<br>**sequence #** |
| --- | --- | --- | --- | --- | --- |
| A_d11 | 10,213,428 | 1,678,431 | 353,052 | 250,000 | 87,520 |
| A_d17 | 4,718,128 | 404,665 | 100,031 | 250,000 | 69,541 |
| A_d45 | 2,446,168 | 215,355 | 99,530 | 215,355 | 99,530 |
| B_d10 | 3,918,963 | 148,132 | 45,698 | 148,132 | 45,698 |
| B_d19 | 3,970,211 | 460,397 | 298,830 | 250,000 | 171,877 |
| C_d6 | 4,206,074 | 538,240 | 310,825 | 250,000 | 157,012 |
| C_d15 | 4,369,267 | 466,795 | 210,434 | 250,000 | 120,680 |
| D_d6 | 4,308,679 | 457,369 | 160,539 | 250,000 | 103,612 |
| D_d28 | 3,593,294 | 142,798 | 84,579 | 142,798 | 84,579 |
| E_d23 | 4,937,896 | 782,329 | 262,323 | 250,000 | 104,363 |
| E_d44 | 3,274,130 | 543,191 | 253,671 | 250,000 | 137,775 |
| E_d99 | 3,900,483 | 276,160 | 58,633 | 250,000 | 54,760 |
| F_d14 | 2,454,273 | 179,398 | 98,942 | 179,398 | 98,942 |
| F_d36 | 2,060,695 | 187,156 | 142,352 | 187,156 | 142,352 |
| G_d9 | 4,698,663 | 626,689 | 223,449 | 250,000 | 104,310 |
| G_d22 | 3,577,375 | 529,997 | 296,335 | 250,000 | 155,254 |

**Supplementary Table 6.** Statistics for the pre-processing of the IGκ and IGλ NGS data.

| Kappa chain (IGκ) repertoire |  |  |  | Lambda chain (IGλ) repertoire |  |  |
| --- | --- | --- | --- | --- | --- | --- |
| Sample | Raw read | UMI-processed read<br>(functional filtering performed) | Unique consensus sequence # | Raw read | UMI-processed read<br>(functional filtering performed) | Unique consensus sequence # |
| A_d11 | 1,147,464 | 8,354 | 2,662 | 2,085,248 | 36,557 | 6,826 |
| A_d17 | 1,916,919 | 19,954 | 5,487 | 1,489,720 | 13,881 | 4,966 |
| A_d45 | 1,315,147 | 46,260 | 19,151 | 1,496,933 | 72,241 | 22,959 |
| B_d10 | 1,298,486 | 18,187 | 6,439 | 961,491 | 12,980 | 1,923 |
| B_d19 | 1,223,146 | 25,509 | 11,159 | 3,590,964 | 357,920 | 56,717 |
| C_d6 | 1,553,360 | 126,170 | 33,153 | 814,108 | 58,115 | 15,939 |
| C_d15 | 1,508,906 | 158,103 | 29,785 | 925,777 | 37,942 | 6,198 |
| D_d6 | 1,628,458 | 44,235 | 19,369 | 1,234,487 | 50,927 | 16,988 |
| D_d28 | 1,189,263 | 81,625 | 25,463 | 1,022,841 | 92,332 | 21,447 |
| E_d23 | 2,916,519 | 58,366 | 12,820 | 1,536,592 | 35,106 | 9,761 |
| E_d44 | 1,634,121 | 64,292 | 19,155 | 1,543,971 | 139,647 | 26,723 |
| E_d99 | 1,224,919 | 30,077 | 13,879 | 1,624,470 | 74,612 | 21,789 |
| F_d14 | 1,439,098 | 8,848 | 2,555 | 1,035,486 | 7,955 | 3,644 |
| F_d36 | 1,340,700 | 62,808 | 21,018 | 889,350 | 48,016 | 13,574 |
| G_d9 | 2,265,376 | 16,048 | 4,147 | 1,310,891 | 8,694 | 4,571 |
| G_d22 | 1,591,445 | 44,963 | 13,327 | 1,028,691 | 16,260 | 5,889 |

**Supplementary Table 7.**
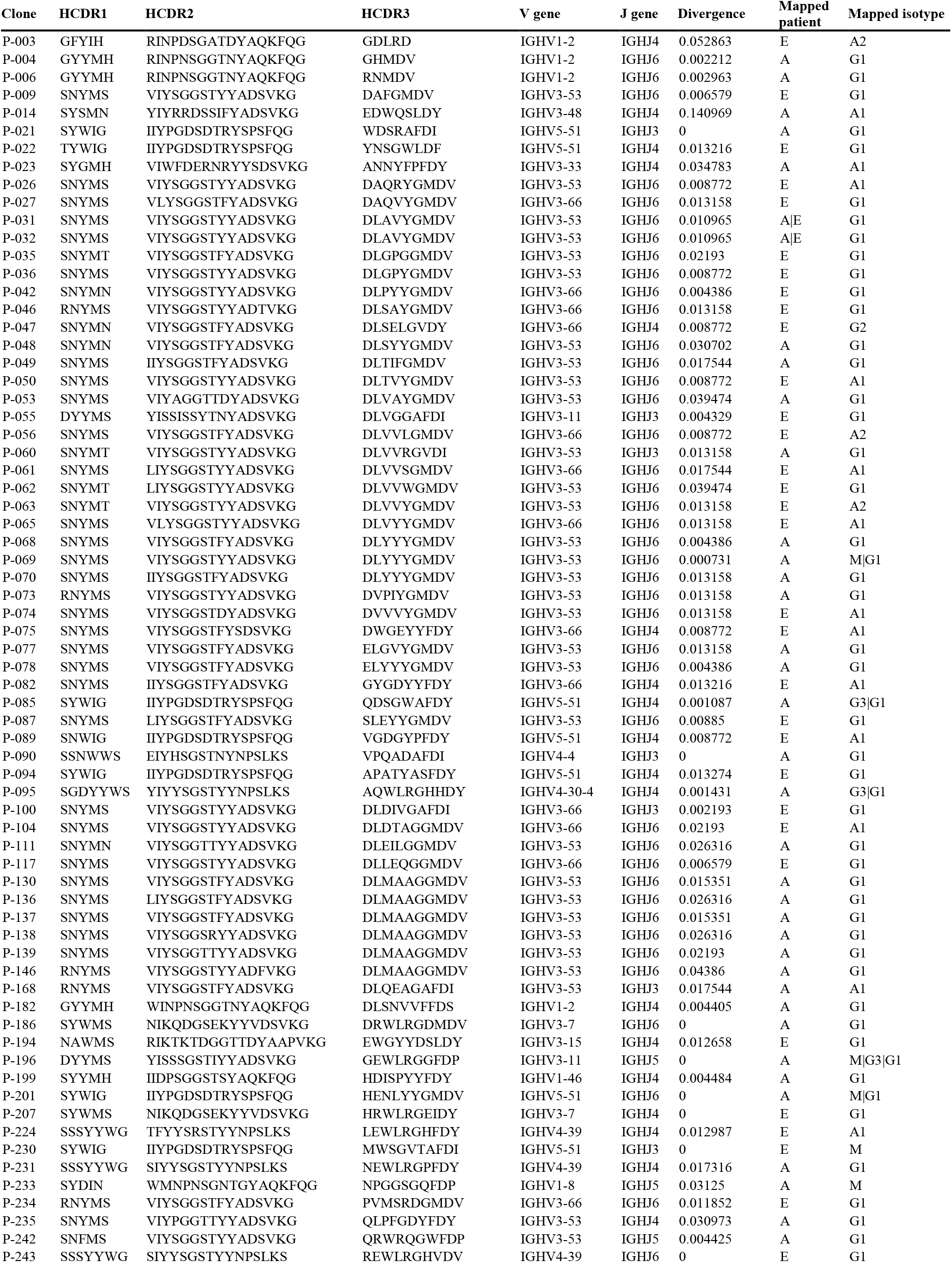
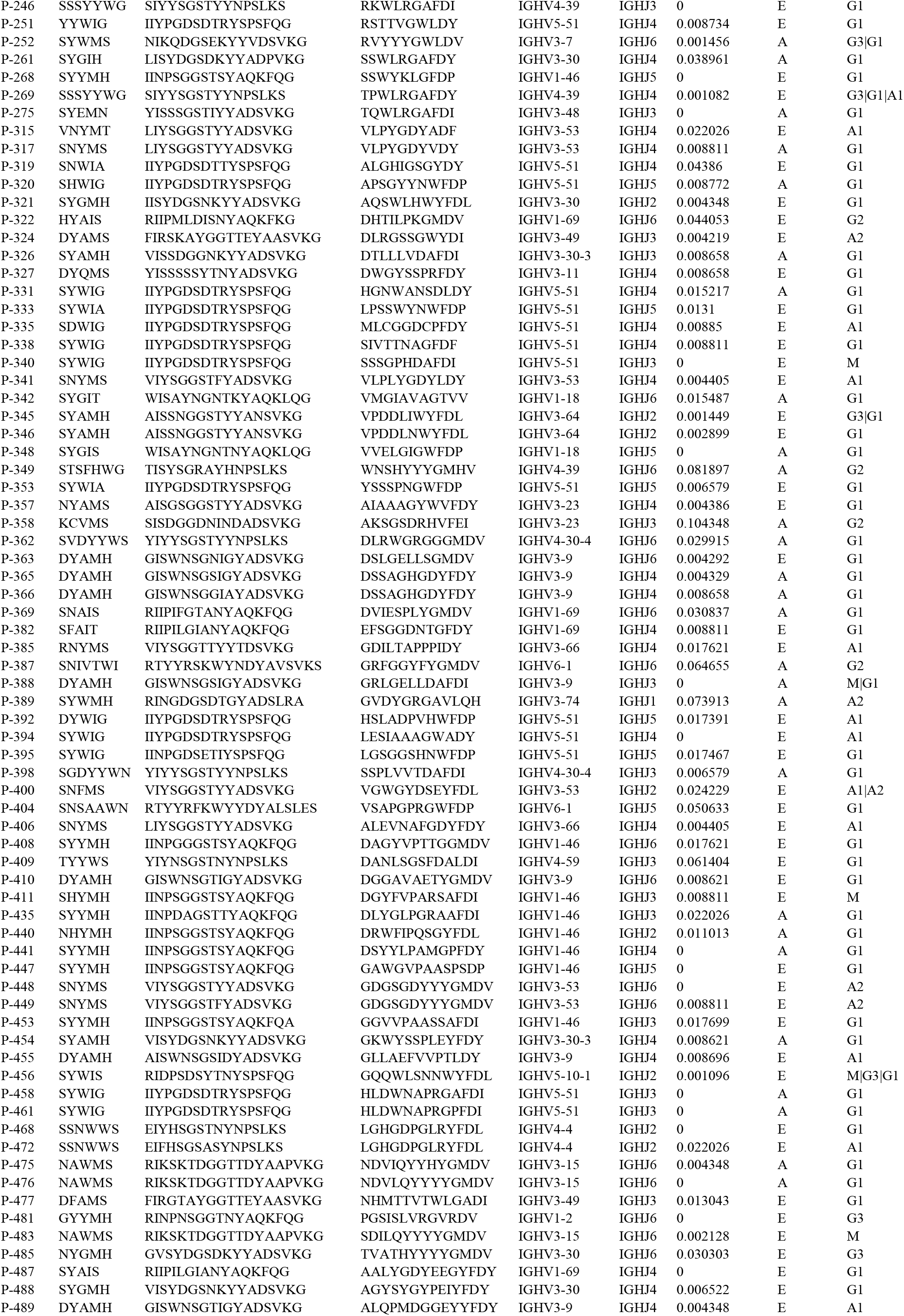
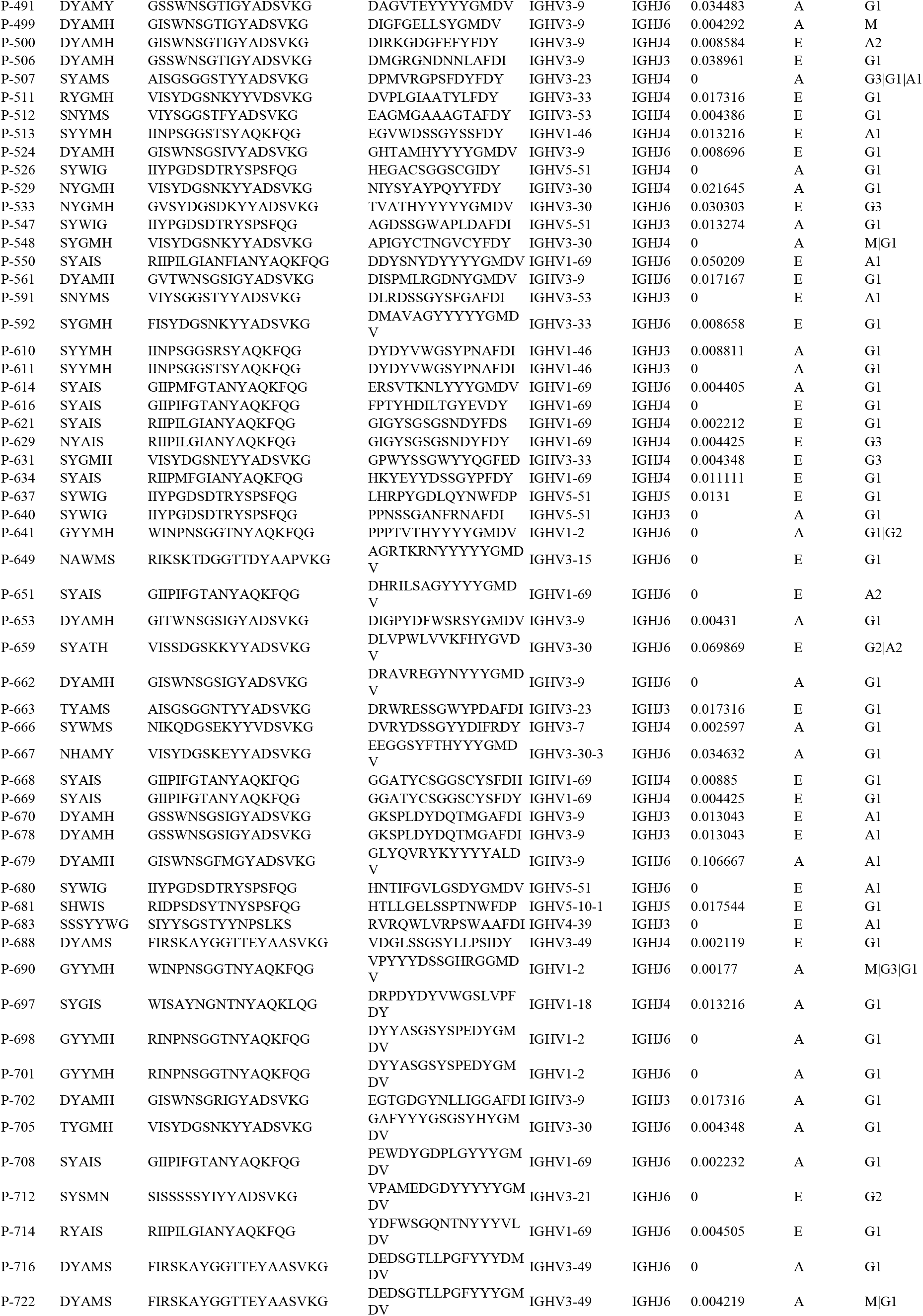
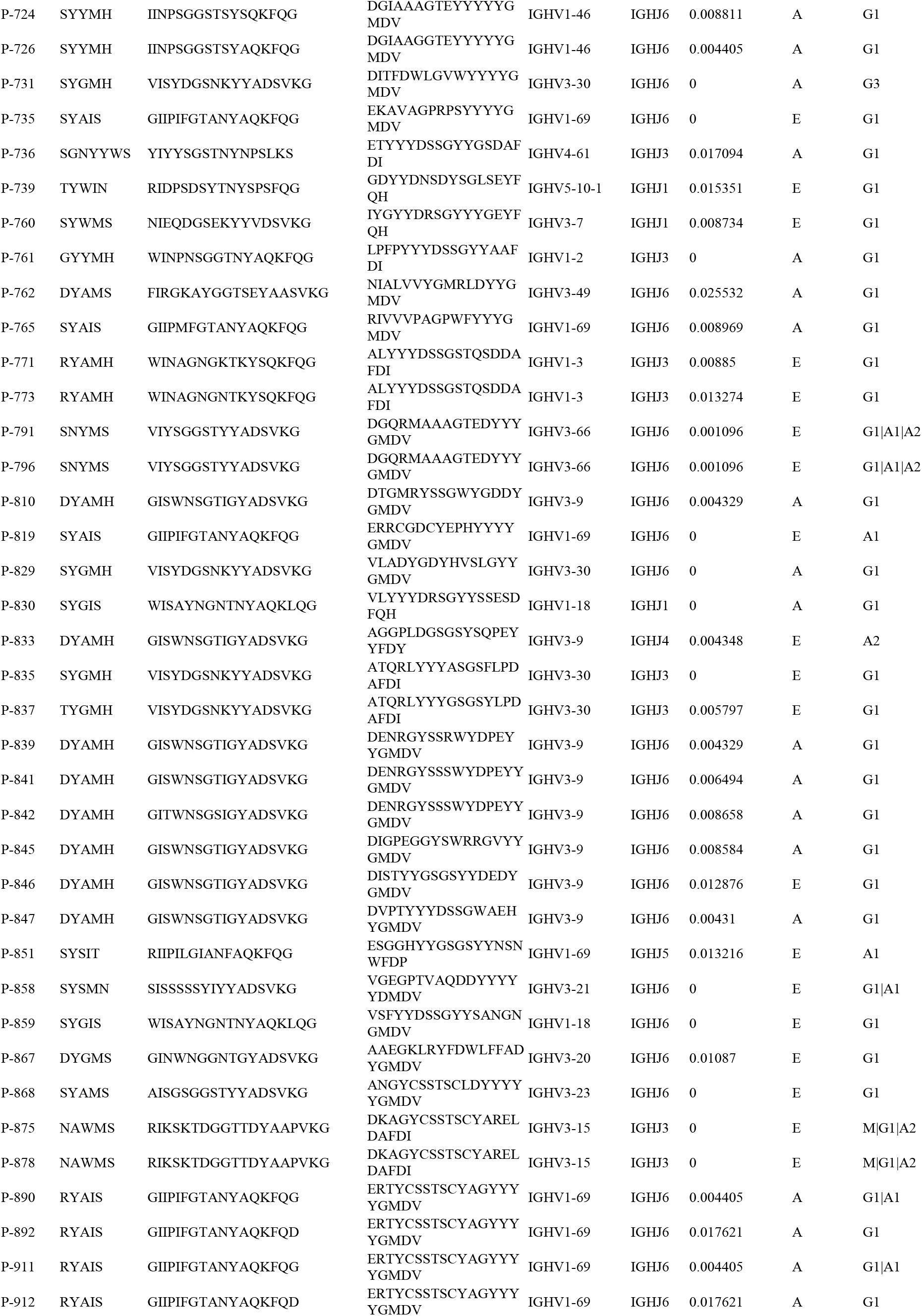
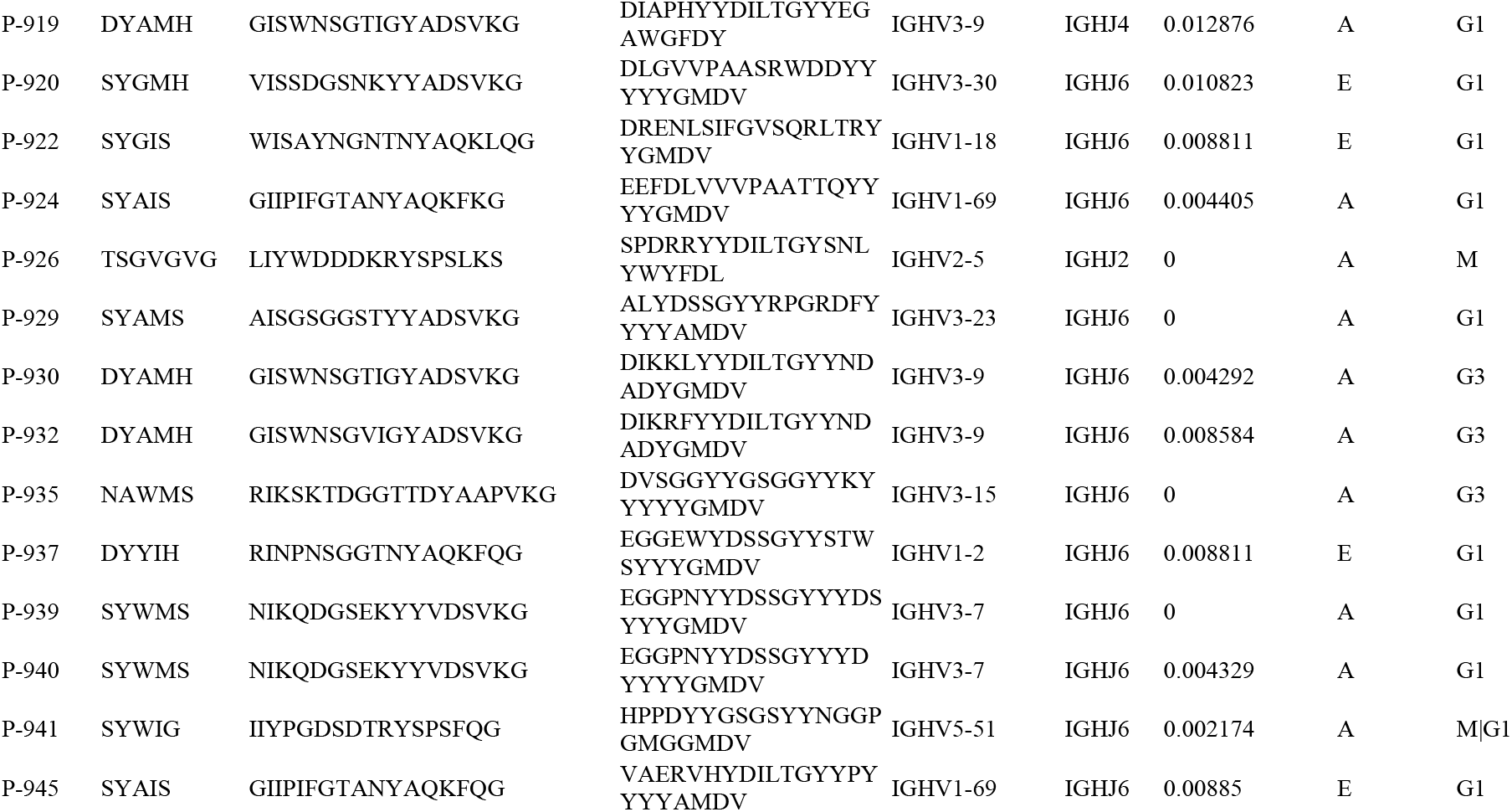
The RBD-binding predicted clones.

**Supplementary Table 8.**
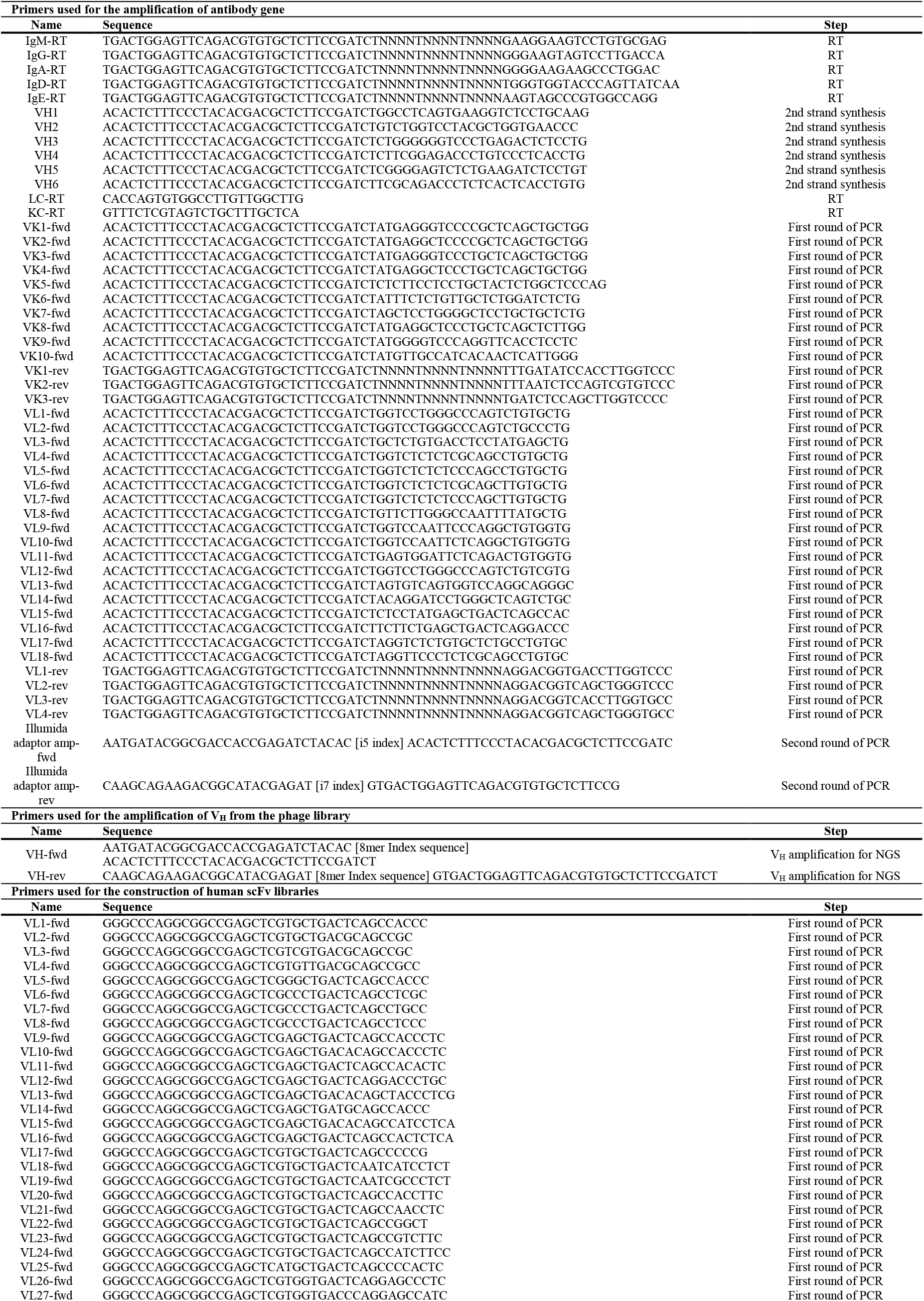
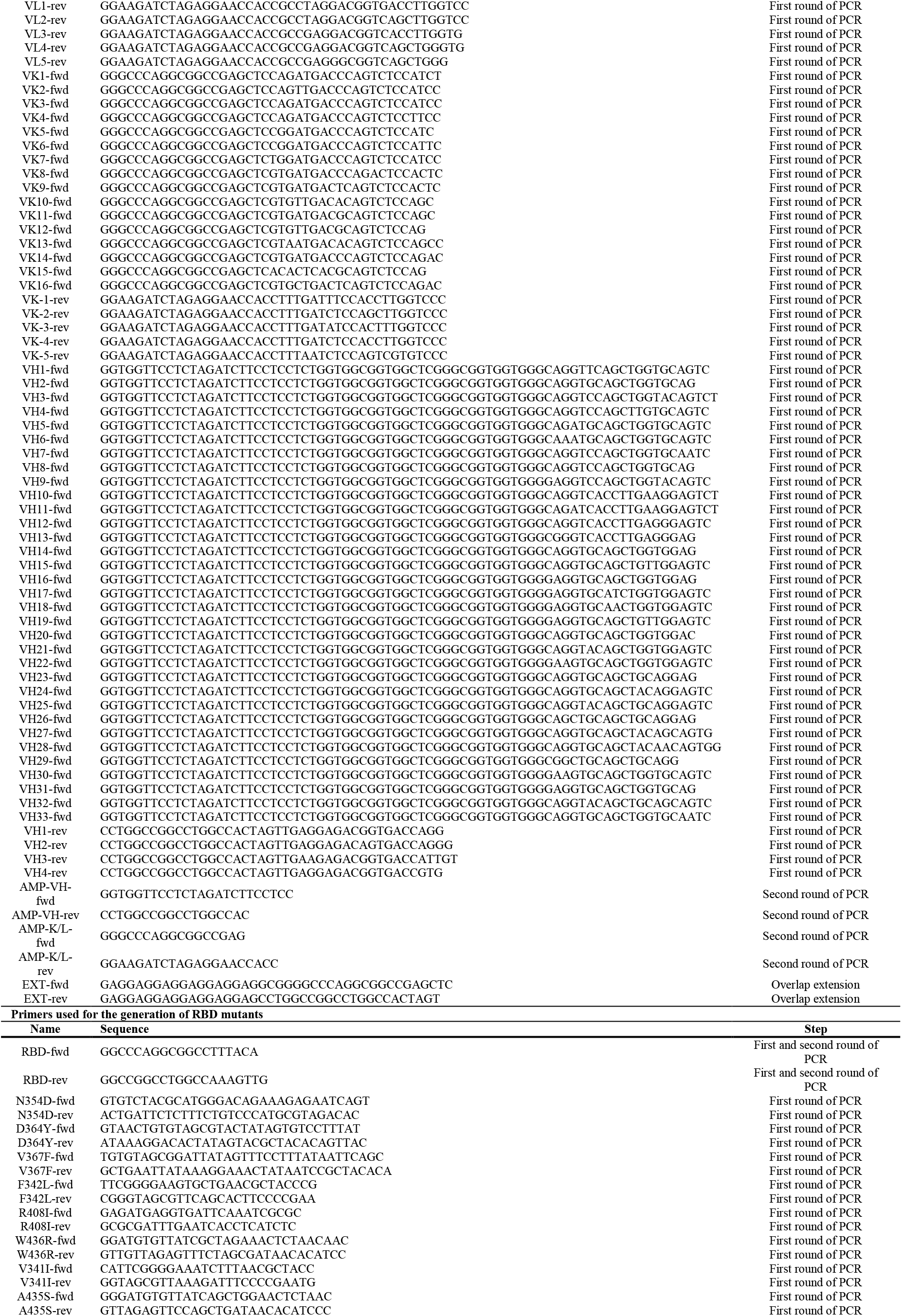
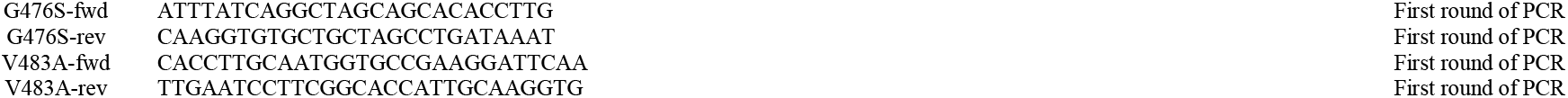
Primers used in the study.

